# Transcriptional dynamics of the murine heart during perinatal development at single-cell resolution

**DOI:** 10.1101/2024.03.05.583423

**Authors:** Lara Feulner, Florian Wünnemann, Jenna Liang, Philipp Hofmann, Marc-Phillip Hitz, Denis Schapiro, Severine Leclerc, Patrick Piet van Vliet, Gregor Andelfinger

## Abstract

Heart maturation and remodelling during the foetal and early postnatal period are critical for proper survival and growth of the foetus, yet our knowledge of the molecular processes involved are lacking for many cardiac cell types. To gain a deeper understanding of the transcriptional dynamics of the heart during the perinatal period, we performed single-cell RNA-seq on E14.5, E16.5, E18.5, P0, P4 and P7 mouse hearts to establish a catalogue of 49,769 cells. Gene regulatory network and pathway activity analyses underscored that heart maturation is strongly associated with regulation of cell growth and proliferation via pathways such as TGFβ. We additionally identified a common, cell type-independent signature for imprinted genes over time. Surprisingly, bioinformatics analyses and confirmation with RNAscope confirmed that while lncRNA H19 expression decreased over time in multiple cardiac cell types, it remained stably expressed in endocardial cells between E14.5 and P7. This suggests a differential requirement for H19 in the endocardium, and points towards an endocardium-specific maturation process when compared to other cardiac cell types. We envision this dataset to serve as a resource for better understanding perinatal heart maturation at the transcriptomic level, and to help bridge the gap between early developmental and adult heart stages for single-cell transcriptomics.

## Introduction

The mammalian heart is a complex, multicellular organ, responsible for blood circulation of deoxygenated blood to the lungs and oxygenated blood to the rest of the body. During early embryonic development, the heart forms from the first and second heart fields to give rise to the four chambered heart, through a series of complex morphogenetic changes and contributions from extracardiac progenitors (Anderson et al., 2014; Bruneau, 2013; Moore-Morris et al., 2018). Heart formation takes place early in embryonic development and major structures are established around seven weeks of gestational life in humans and about 14 days in the mouse. Although the heart is nearly fully formed and functional at this point, it still has to undergo extensive growth and cellular changes throughout foetal and postnatal life (Buckingham et al., 2005; Pervolaraki et al., 2018). The transition from foetal to neonatal life imposes several major challenges on the young heart. Oxygen availability drastically changes, as the foetus progresses from a relatively hypoxic intrauterine environment to atmospheric oxygen levels (Dawson et al., 2010) and the postnatal heart has to adapt to increased workload due to cessation of foetal blood circulation and start of oxygenation via the lungs (Morton and Brodsky, 2016; Tan and Lewandowski, 2020). The large physiological changes imposed on the newborn heart are accompanied by molecular adaptations of the different cell types (Lai et al., 2008; Piquereau and Ventura-Clapier, 2018; Sim et al., 2015). Cardiac muscle cells transition from glycolysis to fatty-acid energy metabolism to meet energy demands (Murphy et al., 2021a, 2021b; Padula et al., 2021; Piquereau and Ventura-Clapier, 2018; Uosaki et al., 2015) and cardiac fibroblasts need to provide structural support via extracellular matrix (ECM) synthesis (Hortells et al., 2020; Wang et al., 2020). Several studies have investigated heart maturation using bulk technologies such as RNA-seq and ATAC-seq, often with a focus on cardiomyocytes (Quaife-Ryan et al., 2017; Talman et al., 2018). Although these efforts have uncovered important maturation signals for cardiomyocytes, such as oestrogen-related receptor signalling (Sakamoto et al., 2020), non-myocyte cell populations are often neglected.

Single cell RNA-seq has become a powerful tool to interrogate the transcriptome of the different cell types of the heart (Paik et al., 2020; Yamada and Nomura, 2020). Early studies investigating heart development in the mouse discovered important transcriptomic changes across major cardiac cell types, but were limited in throughput and the number of different cell types identified (DeLaughter et al., 2016; Li et al., 2016). Subsequent technological advances have enabled much higher throughput for mouse and human heart single-cell transcriptomes to study cardiac cell diversity and disease models (Asp et al., 2019; Cui et al., 2019; Litviňuková et al., 2020; Skelly et al., 2018; Tucker et al., 2020). Despite these improvements, most single-cell studies focused on models of early developmental processes in heart formation (de Soysa et al., 2019), postnatal cardiomyocyte maturation and regeneration (Hu et al., 2018; Kannan et al., 2021; Murphy et al., 2021b; Wang et al., 2020) or acute disease models like myocardial infarction and cardiomyopathy (Forte et al., 2020; Koenig et al., 2022; Li et al., 2019; Rizzo et al., 2023; Ruiz-Villalba et al., 2020; Tombor et al., 2021; Vafadarnejad et al., 2020). This has left a gap in our understanding of the transcriptomic changes in the heart at the single-cell level during the perinatal window, the time just before and right after birth, which is a critical period for heart growth. Few studies have covered this period, with Feng et al. (Feng et al., 2022) being a recent exception.

Here, we generated single-cell transcriptomes from mouse hearts across six time points, during foetal to neonatal transition (further referred to as the perinatal period, time points E14.5 to P7) using the droplet-based technology Drop-seq (Macosko et al., 2015). We identified distinct cardiac cell types and states and investigated the cell-type specific and common temporal gene expression dynamics. We performed gene regulatory network and pathway activity analysis to infer specific transcriptional programs associated with heart maturation. We examined gene expression changes over the perinatal period and identified the lncRNA H19 as significantly decreasing over time in multiple cell types, except in endocardial cells. Our data, results and insights reveal transcriptional dynamics during the perinatal period in all major cardiac cell types and can serve as guidance to identify new potential targets for cell maturation in *in vitro* cardiac differentiation protocols.

## Results

### Single-cell diversity in the mouse heart during the perinatal maturation

To establish a dataset of cardiac cell types throughout the perinatal period in the mouse, we performed scRNA-seq using Drop-seq on dissociated whole mouse hearts at three foetal (E14.5, E16.5, E18.5) and three neonatal (P0, P4, P7) time points. After quality control, we obtained a final merged dataset of 49,769 single-cell transcriptomes (E14.5: 8,889 cells, E16.5: 8,396 cells, E18.5: 6,058 cells, P0: 9,213 cells, P4: 8,922 cells, P7: 8,291 cells) across perinatal heart remodelling and maturation stages (**Figure 1, Table S1**). Seurat clustering and manual annotation using known markers revealed common cardiac cell types across the six sampled time points (**Figure 2, Table S2**).

**Figure 1:**
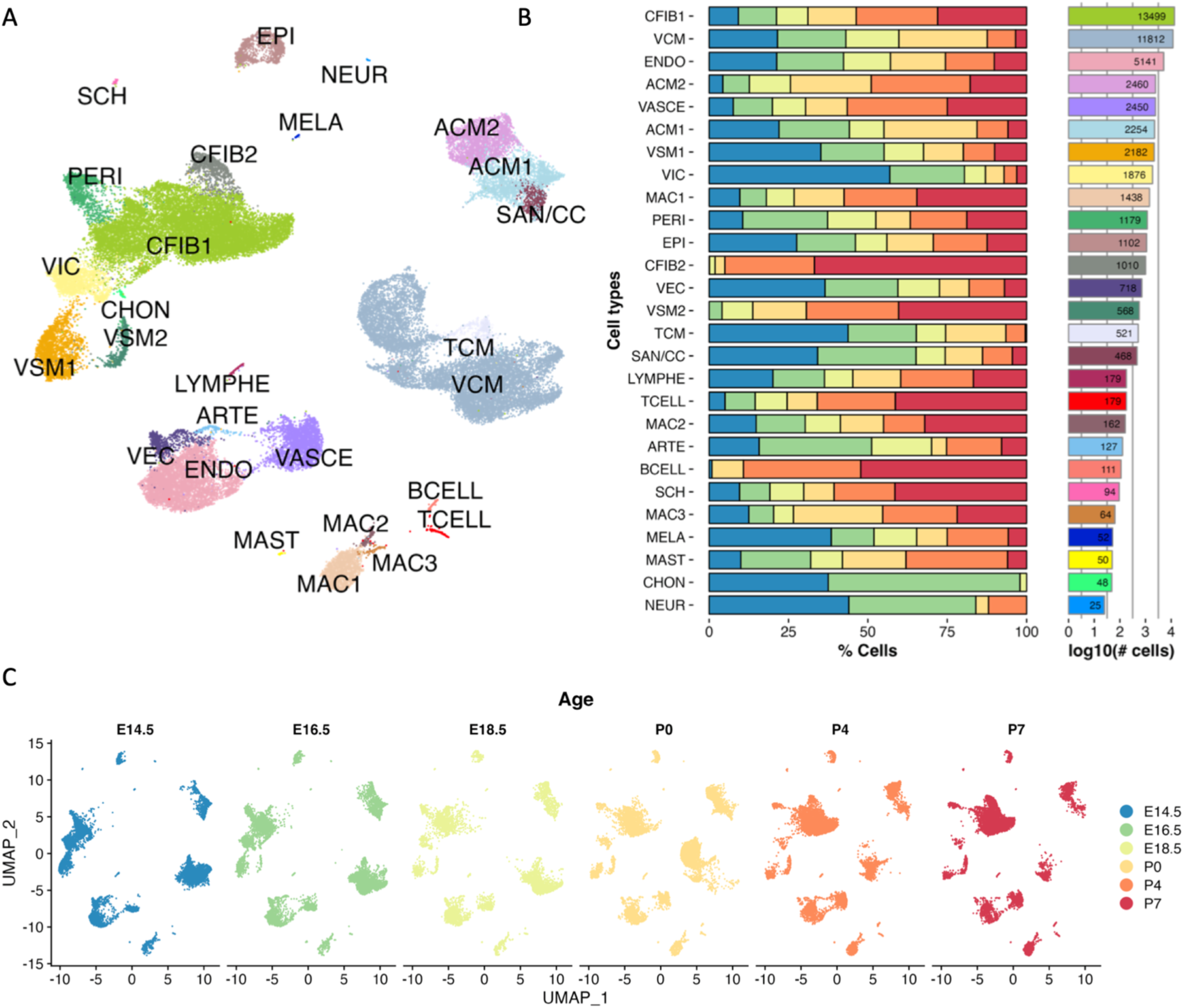
**A)** Two dimensional UMAP embedding of 49,769 cardiac cells from E14.5, E16.5, E18.5, P0, P4 and P7 mice. Six replicates were used per timepoint for a total of 36 different samples. **B)** The fraction of cells across cell types and time. Cell types are ordered by the total number of cells from top to bottom. Numbers in the right-side bars represent total cell count, while x-axis is on a log10 scale. **C)** UMAP embedding plots for individual timepoints. Known biological progression can be observed, for example the lack of fibroblasts at E14.5 with extensive proliferation thereafter.

**Figure 2:**
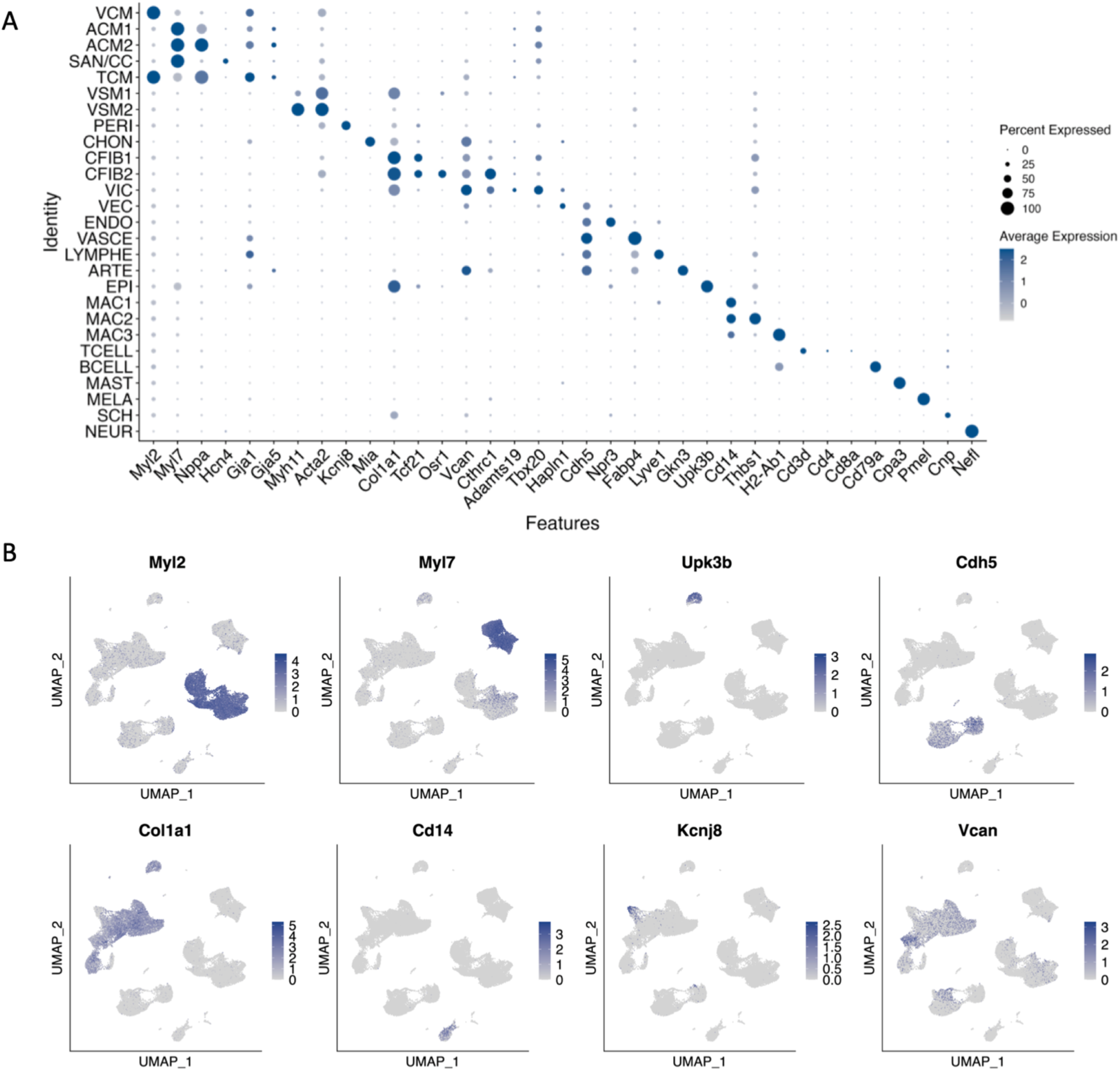
A) Dotplot of marker genes for specific cardiac cell types used to guide cluster annotation. B) Feature plots of selected marker genes (*Myl2*: VCM, *Myl7*: ACM, *Upk3b*: EPI, *Cdh5*: Endothelial, *Col1a1*: CFIB, *Cd14*: MAC, *Kcnj8*: PERI, *Vcan*: VIC).

Cardiomyocytes (CMs) were an abundant cell type in our dataset (35% of all cells), consistent with relative cell quantities in the heart (Banerjee et al., 2007; Pinto et al., 2016; Zhou and Pu, 2016). Three major cardiomyocyte subtypes, atrial (ACM: *Myl7*), ventricular (VCM: *Myl2, Myl3, Myh7*) and trabecular (TCM: *Nppa*, *Gtsf1*) were identified. Trabecular cardiomyocytes mostly resembled ventricular cardiomyocytes in their transcriptional signature, but share several known markers with atrial cardiomyocytes, such as expression of atrial natriuretic peptide A (*Nppa*) (Houweling et al., 2005; Seidman et al., 1991). Two atrial cardiomyocyte subtypes (ACM1 and ACM2) were noted, primarily distinguished by increased expression of *Nppa*, *Nppb,* and *Bmp10* in ACM2. High-resolution clustering additionally revealed an ACM1-like, *Nppa*-negative cluster of cardiomyocytes expressing genes associated with the cardiac conduction system such as *Vsnl1*, *Shox2,* and *Slc22a1* (Goodyer et al., 2019), likely containing cells of the sinoatrial node and the rest of the cardiac conduction system (SAN/CC).

Mural and interstitial cells represented the largest group of cells that were readily sampled at all time points (41% of cells). These cells were comprised of cardiac fibroblasts (CFIB: *Dpt*, *Tcf21*, *Lsp1, Spon2*), vascular smooth muscle cells (VSM: *Tagln*, *Acta2*, *Rgs5*), pericytes (PERI: *Kcnj8*, *Abcc9*, *Higd1b*) and valvular interstitial cells (VIC: *Vcan*, *Col9a1, Prss35, Adamts19, Cthrc1*). Both CFIB and VSM were subdivided in two clusters, with CFIB2 being notably enriched for *Osr1* and VSM2 being enriched for the contractile gene *Myh11*. In contrast to the interstitial and mural cell populations which shared some similarity in their transcriptional profiles, epicardial cells (EPI) expressed highly specific genes (*Upk3b*, *Upk1b*, *Wt1*, *Msln, Krt19*).

Several different subtypes of endothelial cells (ECs, 17% of cells) were additionally present in our dataset. ECs were generally identifiable by canonical markers VE-Cadherin (*Cdh5)* and CD31 (*Pecam1),* and further subdivided into vascular (VASCE: *Fabp4*, *Cav1*), endocardial (ENDO: *Npr3*, *Mmrn2, Emcn),* lymphatic (LYMPHE: *Ccl21a*, *Mmrn1*) and valvular endothelial cells (VEC: *Hapln1*, *Cldn11*, *Lsamp*, *Cd24a*). An additional endothelial cluster was classified as arterial (ARTE) based on strong expression of *Gkn3* (Vanlandewijck et al., 2018) in addition to *Gja4*, *Fbln5*, *Bmx,* and *Eln*.

Macrophages were the predominant immune cell subtype found among cardiac cells. We identified three Cd14+ macrophage subsets: MAC1 (*C1qa*, *Mrc1*, *Pf4*, *Ccl2*), MAC2 (*Plac8*, *S100a8*, *Thbs1*) and MAC3 (*H2-Eb1*, *H2-Aa*, *H2-Ab1*, *Cd74*). Other myeloid and lymphoid cell types were seen at later developmental stages representing B-cells (BCELL: *Cd79a*, *Ighm*, *Ly6d*), T-cells (TCELL: *Trbc2*, *Ms4a4b*, *Cd3d*) and Mast cells (MAST: *Cpa3*, *Hdc*, *Il1rl1*).

Finally, we identified small but clearly distinct clusters of melanocytes (MELA: *Pmel*, *Mlana*, *Dct*, *Plp1*), Schwann cells (SCH: *Erbb3*, *Mal*, *S100b*, *Plp1*) and neuronal cells (NEURO: *Nefl*, *Nefm*, *Elavl2*). Lastly, a small cluster of E14.5-E16.5 cells enriched for *Col2a1*, *Mia,* and *Wif1* was identified as probable chondrocytes (CHON), and thus likely of tracheal rather than cardiac origin. These cells were therefore excluded from subsequent analysis. While most cell types were found in all biological replicates at all time points, rare cell types such as neuronal cells, melanocytes, B- and T cells were only sampled in some of the biological replicates we used, owing to their low relative abundance in each isolated heart.

To experimentally validate the cell specificity of our markers, we performed RNAscope using probes for *Myl2*, *Myl7*, *Upk3b*, *Tcf21*, *Myh11*, *Kcnj8*, *Fabp4*, *Npr3*, *Cdh5,* and *Cthrc1* (**Figure 3**). VCM, ACM, and EPI cells were especially well stained by *Myl2*, *Myl7,* and *Upk3b* respectively. While no truly unique markers (i.e., not shared by other mesenchymal cell types) covering all time points could be found for VICs, *Cthrc1* was fairly specific to valves up to P0, after which significant *Cthrc1* expression was observed in non-VIC fibroblasts. We were particularly interested in *Npr3*, which prior studies had found to be an endocardial marker (Feng et al., 2019; Zhang et al., 2016). Our RNAscope showed *Npr3* to specifically outline atrial pectinate and ventricle trabecular endocardial cells without staining VECs, lending credence to *Npr3* being a strong marker for non-valve chamber endocardium. Overall, the RNAscope staining validated the markers used for annotating our scRNA-seq clusters.

**Figure 3:**
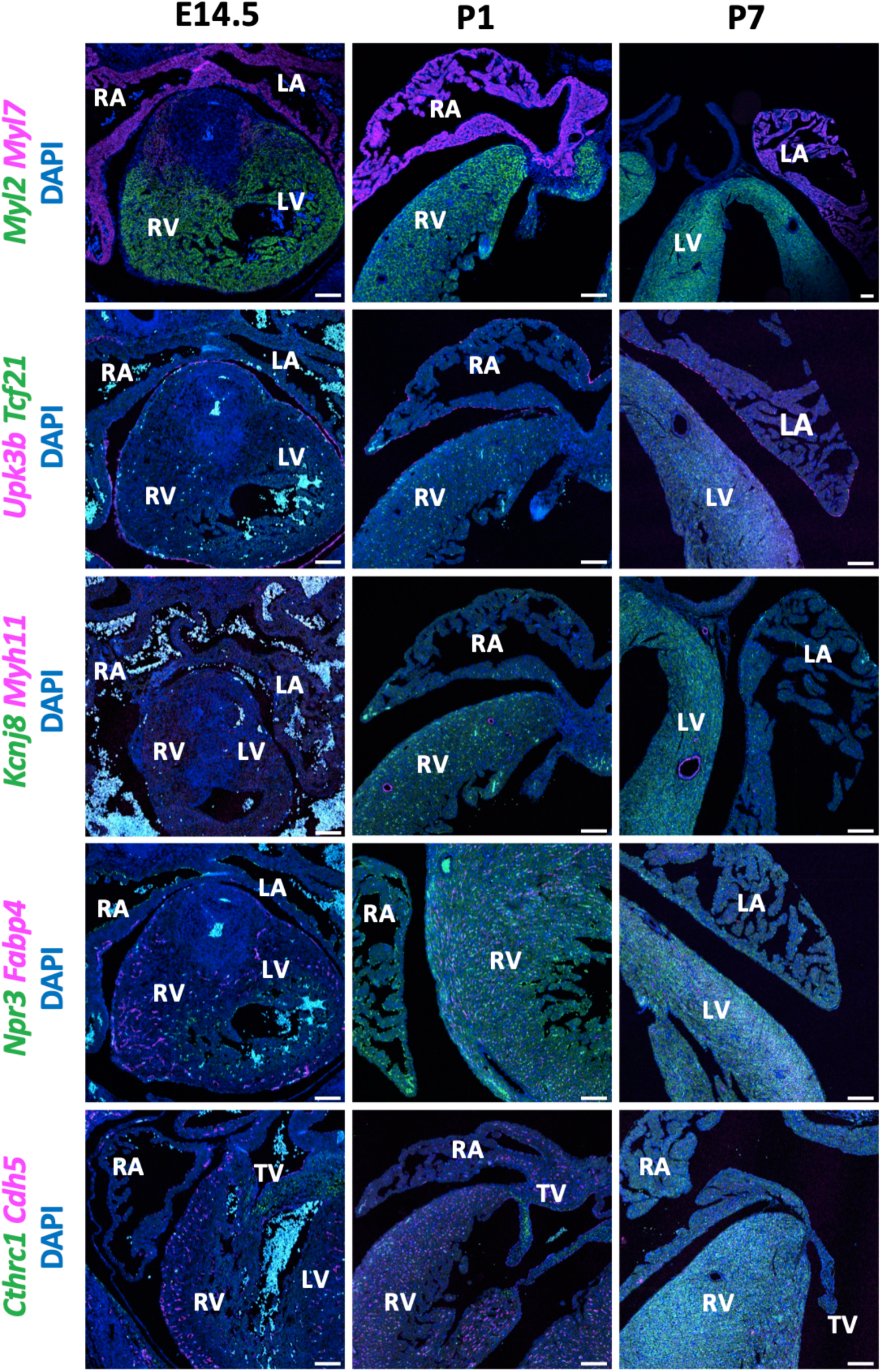
RNAscope stains for selected marker genes across E14.5, P1 and P7 timepoints. *Myl2* and *Myl7* stain for ventricular cardiomyocytes and atrial cardiomyocytes respectively. *Tcf21* and *Upk3b* stain for cardiac fibroblast and epicardial cells. *Kcnj8* and *Myh11* stain for pericyte and vascular smooth muscle. *Npr3* and *Fabp4* stain for endocardial cells and vascular endothelial cells. Lastly, *Cthrc1* and *Cdh5* stain for valve interstitial cells and endothelial cells.

### Gene regulatory network and pathway activity analyses underscore common importance of cell proliferation regulation across cardiac cell types

Following identification of distinct cell types in our dataset, we used the tool decoupleR (Badia-I-Mompel et al., 2022) to analyse gene regulatory network and pathway activity changes across developmental timepoints. Cell types with sufficient population and representation from E14.5-P7 were each isolated from the main dataset and analysed separately. Altogether we obtained separate results for four cardiomyocyte populations (ACM, VCM, TCM, SAN/CC), five endothelial populations (ENDO, VASCE, VEC, LYMPHE, ARTE), cardiac fibroblasts, epicardial cells, valve interstitial cells, vascular smooth muscle, pericytes, and macrophages.

### Gene regulatory network analysis

The decoupleR package was used to infer transcription factor (TF) activity scores from a curated collection of TFs and their transcriptional targets (CollectTRI, see Methods). For each cell in a dataset representing a single cardiac cell type, a positive score indicated a TF was active, a negative score indicated the TF was inactive and 0 indicated lack of regulation by the TF. The mean activity scores for each timepoint were then calculated and standard deviation was used to assess which TFs showed the greatest variation in scores across time. The mean scores per time point for these top TFs were then plotted as a heatmap, with red and blue representing increased and decreased TF activity respectively (**Figure 4a, Figure S1**). A notable shift in TF activity was found to occur from 18.5 to P0 (i.e. pre- to postnatal) in most cell types, indicating that postnatal maturation in those cells is associated with significant changes in molecular pathways requiring the activation of distinct TFs.

**Figure 4:**
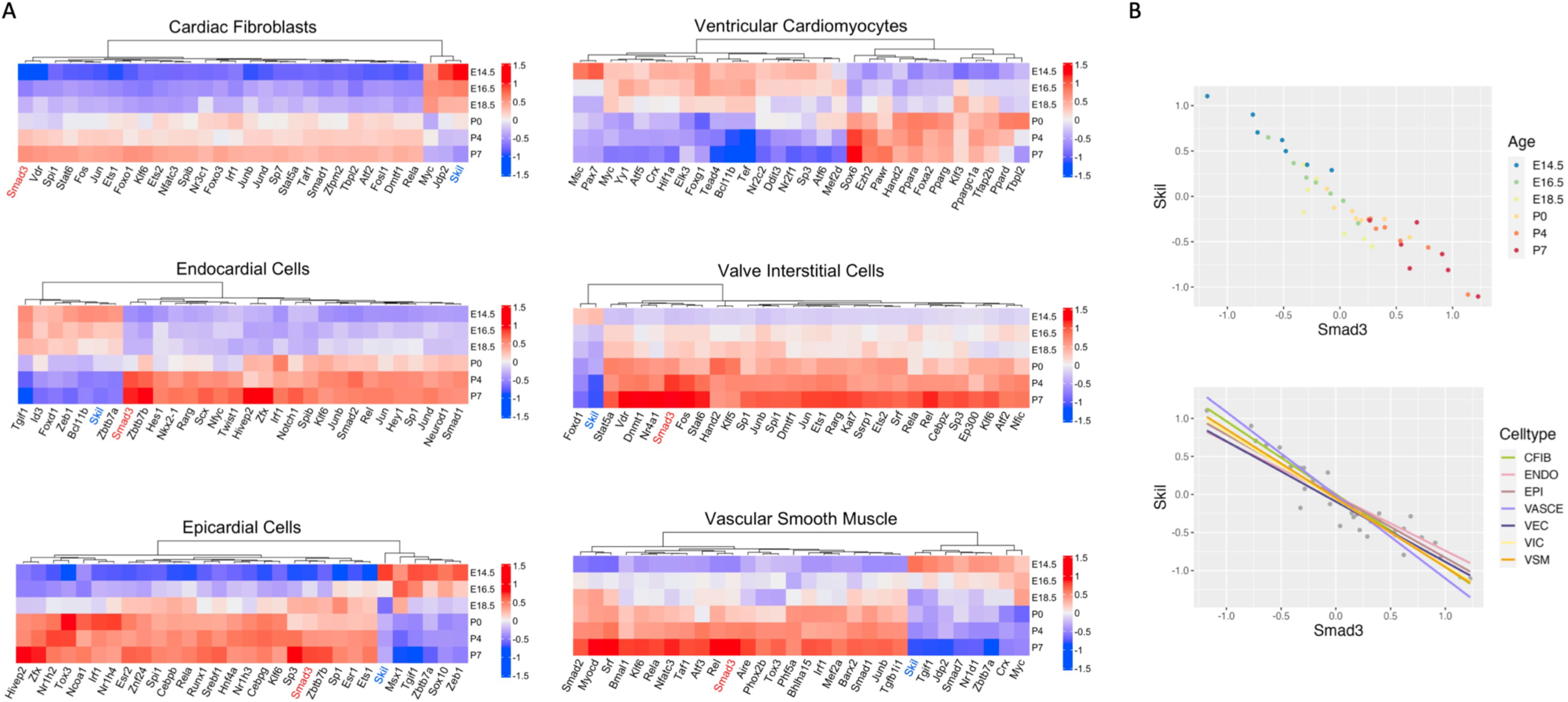
**A)** Transcription factor (TF) activity heatmaps for fibroblasts, ventricular cardiomyocytes, endocardial cells, valve interstitial cells, epicardial cells, and vascular smooth muscle cells. Numbers in legend represent the mean TF activity scores inferred by decoupleR for each time point in that cell type, with positive scores (red) indicating increased activity and negative scores (blue) indicating decreased activity. The TFs shown are those with the greatest variation in activity scores across time per standard deviation analysis. A foetal-postnatal shift in TF activity can be seen, with this transition occurring notably earlier in VICs. Note the prevalence of Skil (blue, downregulated with age) and Smad3 (red, upregulated with age) in multiple cell types. **B)** Scatter plots showing an inverse relationship between Skil activity and Smad3 activity levels, consistent with Skil being a direct inhibitor of Smad2/3 and thus the TGFβ pathway.

Interestingly, this transition occurred much earlier in VICs, consistent with their relatively early initiation and completion of development (E9.0 to E14.5) compared to other cardiac cell types. In many cases higher postnatal activity predominated among the top TFs (with fibroblasts and VICs being extreme examples), but cardiomyocyte, macrophage, and pericyte TFs were more evenly split between foetal and postnatal activity. Arterial endothelium and trabecular cardiomyocytes showed particularly extreme change in TF activity from P4 to P7.

Overall, the top TFs were less cell type specific and more general than anticipated. Two proto-oncogenes, *Myc* and *Skil*, were repeatedly detected among the top hits showing significantly decreased activity across time. *Skil* was particularly interesting as Smad2-3, members of the TGFβ pathway and known to be inhibited by Skil (Luo et al., 1999), were also often among the top hits with increased activity across time. Correlation analysis of *Skil* vs *Smad3* activity indeed showed a clear strong negative correlation between the two transcription factors (**Figure 4b**). The balance between *Skil* and *Smad2-3* in turn likely plays a role in regulating TGFβ.

Other common TFs directly regulating growth and proliferation included *Fos/Jun* family members, which were among the top results in fibroblasts, VICs, vascular smooth muscle, and endothelial cells. Finally, peroxisome proliferators including *Ppara/g* were identified by decoupleR in pericytes, vascular endothelium, vascular smooth muscle, and all cardiomyocytes; the latter consistent with their known shift to fatty acid oxidation. In addition to regulating metabolism, PPARγ inhibits cell proliferation via decrease in c-Myc, whereas PPARα induces apoptosis (Martinasso et al., 2007).

Parallel to using decoupleR, we performed gene regulatory network analysis using SCENIC (Aibar et al., 2017; Van de Sande et al., 2020). Briefly, regulon specificity scores (RSS) for both foetal and postnatal timepoints were calculated for each cell type. The resulting scores were then placed in descending order to obtain the top foetal and postnatal TFs (**Table S3**). Foetal timepoints showed some enrichment for TFs known to be expressed or significantly associated with their respective cell type development, ex. *Nkx2-5* and *Hand1/2* in cardiomyocytes (Lints et al., 1993; McFadden et al., 2005), *Etv2* in endocardial cells (Ferdous et al., 2009), *Isl1* in arterial endothelium (Cai et al., 2003; Sun et al., 2007) and *Hes1* in epicardium (del Monte et al., 2011). While *Wt1* was not among the top TFs in epicardial cells, it was enriched at foetal timepoints in a number of other cell types including arterial endothelium, fibroblasts, vascular smooth muscle, and cardiomyocytes, consistent with Wt1 expression in non-epicardial cells at later developmental stages (Rudat and Kispert, 2012). Postnatal time points were commonly enriched for *Fos and Jun* as well as *Klf* factors, particularly *Klf2* and *Klf9*. cJun regulates cell cycle progression and apoptosis and is required for normal heart development (Eferl et al., 1999; Zhang et al., 2013), while Klf2 is a key shear stress response agent required for normal valve development (Chiplunkar et al., 2013; Goddard et al., 2017). Taken together it appears that cell-specific GRNs are more likely to be active during lineage specification early in development (prior to the majority of time points studied here), while heart maturation is a more common and non-cell specific process. This would be consistent with Cui et al.’s finding that cardiomyocytes from 5-7 week human embryos were enriched for *HAND1/2*, *NKX2-5*, and *TBX5* and already had chamber-specific expression profiles (Cui et al., 2019).

### Pathway analysis

Pathway analysis was also performed in decoupleR using the Hallmark geneset collection from the Molecular Signatures Database (MSigDB, (Castanza et al., 2023; Subramanian et al., 2005). Similarly to the GRN analysis, for each cell, an overrepresentation score for each Hallmark pathway was inferred (see Methods). The mean score per time point was then calculated and standard deviation used to assess which pathways showed the most variation across time. Heatmap plots of the mean overrepresentation scores per time point of these top pathways for each respective cell type were then generated (**Figure 5, Figure S2**).

**Figure 5:**
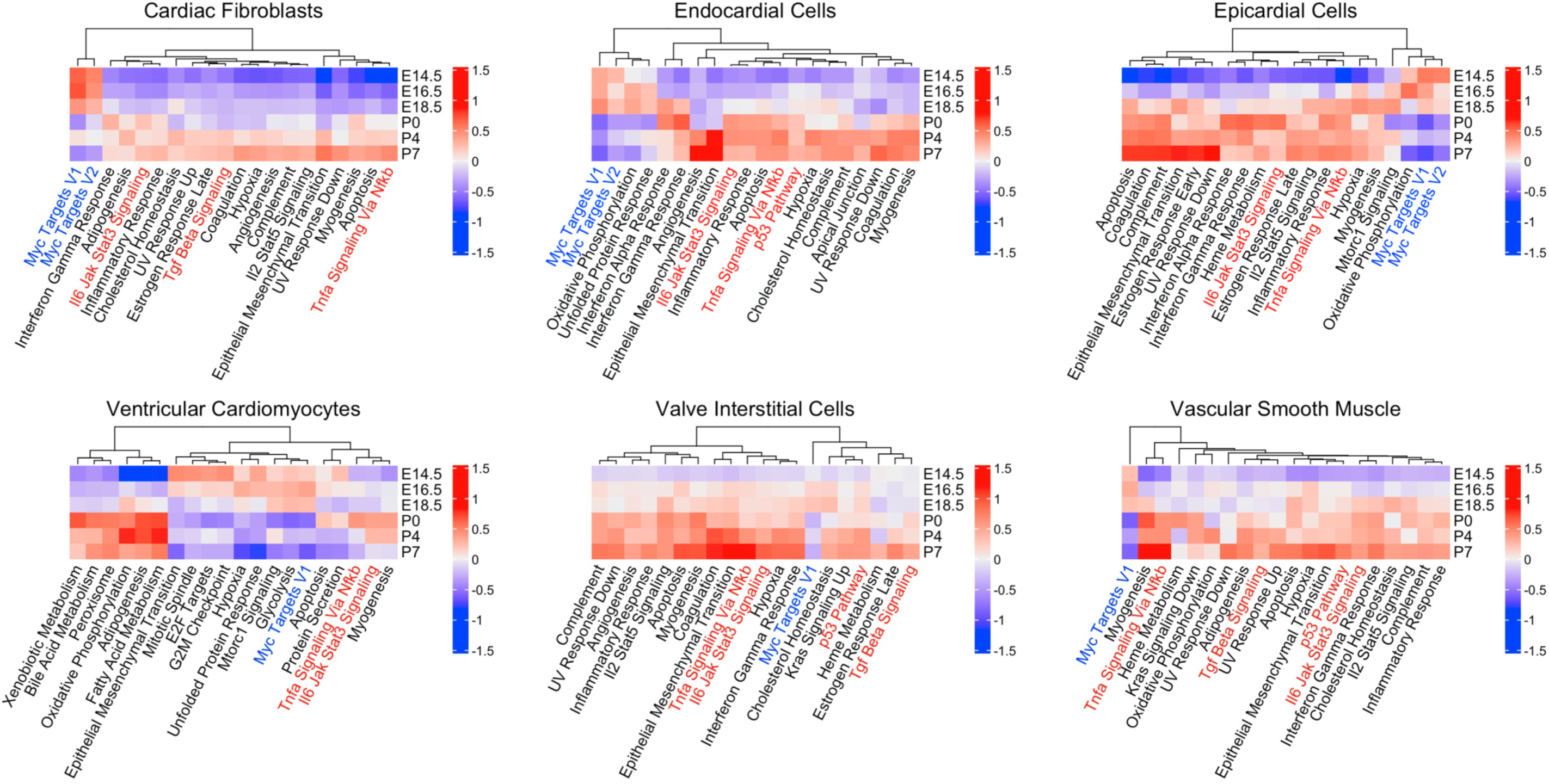
Hallmark pathway activity heatmaps for fibroblasts, ventricular cardiomyocytes, endocardial cells, valve interstitial cells, epicardial cells, and vascular smooth muscle cells. Numbers in legend represent the mean pathway overrepresentation scores inferred by decoupleR for each time point in that cell type, with positive scores (red) indicating increased pathway activity and negative scores (blue) indicating decreased activity. The pathways shown are those with the greatest variation in scores across time per standard deviation analysis. Similar foetal-postnatal shifts as noted from the TF activity analysis can be seen. Note the increased oxidative phosphorylation and fatty acid metabolism over time in VCMs, consistent with known postnatal changes. The Myc pathway (blue) commonly decreases with time while TGFβ, Stat3, TNFα and p53 pathways (red) increase.

A shift in pathway activity was again noted as cell types progressed in development, particularly going from foetal to postnatal (with VICs again showing an earlier shift). It’s worth noting that the pathways did not always show straight increases or decreases from E14.5 to P7. In fact, in many cases a “reversal” from P0 to P4 or P4 to P7 occurred. Myc and the Pi3K-Mtor pathway were the main pathways commonly downregulated with developmental time. Upregulated pathways included Tnfa via Nfkb, IL6 Jak-Stat3, and the p53 and TGFβ pathways. These are known to be associated with hypertrophic cell growth (Dobaczewski et al., 2011; Kodama et al., 1997; Mak et al., 2017; Purcell et al., 2001) and apoptosis (Dhingra et al., 2009) in cardiomyocytes. The TGFβ pathway was additionally found upregulated with time across multiple cell types when using the PROGENy dataset (Schubert et al., 2018) (**Figure S3**), lending further credence to the previous TF activity analysis results. While the referenced studies involved cardiomyocytes, our results would suggest that regulation of cell growth is an important aspect of maturation across all cardiac cell types. Evidence of the known switch from glycolysis to aerobic respiration and fatty acid metabolism in cardiomyocytes (Padula et al., 2021; Uscategui Calderon et al., 2023) could be seen in the results for ACM, VCM, and TCM, with the former being downregulated and the latter two upregulated with time.

### Cell-type specific gene expression dynamics during cardiac foetal to postnatal transition

We next aimed to determine the genes involved in driving heart maturation during perinatal development. We utilised a semi-supervised approach termed psupertime (Macnair et al., 2022), which used known time-series labels (in our case E14.5, E16.5, E18.5, P0, P4, P7) as input to assign cells a quantitative pseudotime score per cell type. The use of pseudotime was important, as not all cells progress through development at the same rate. An ordinal logistic regression model identified genes for each cell type showing differential expression across the pseudotime trajectories during foetal to neonatal transition. Any gene with a non-zero beta coefficient was considered by psupertime to be relevant to the pseudotime ordering of the cells, and thus significantly vary in expression with time (also referred to as showing “dynamic” expression).

Our analysis identified known isoform switches from alpha- (*Myh6*) to beta-myosin heavy chains (*Myh7*) as well as the switch from foetal to adult Troponin I genes (*Tnni1* to *Tnni3*) in ventricular cardiomyocytes, in line with previous observations (Bedada et al., 2014; DeLaughter et al., 2016; Siedner et al., 2003) (**Figure 6a**). These positive control genes indicate that our approach is able to confidently identify known, biologically relevant signals with temporal dynamic expression. We applied the same strategy to all major cell types, excluding those with inconsistent sampling across time or very few total cells in the dataset (**Table S4**).

**Figure 6:**
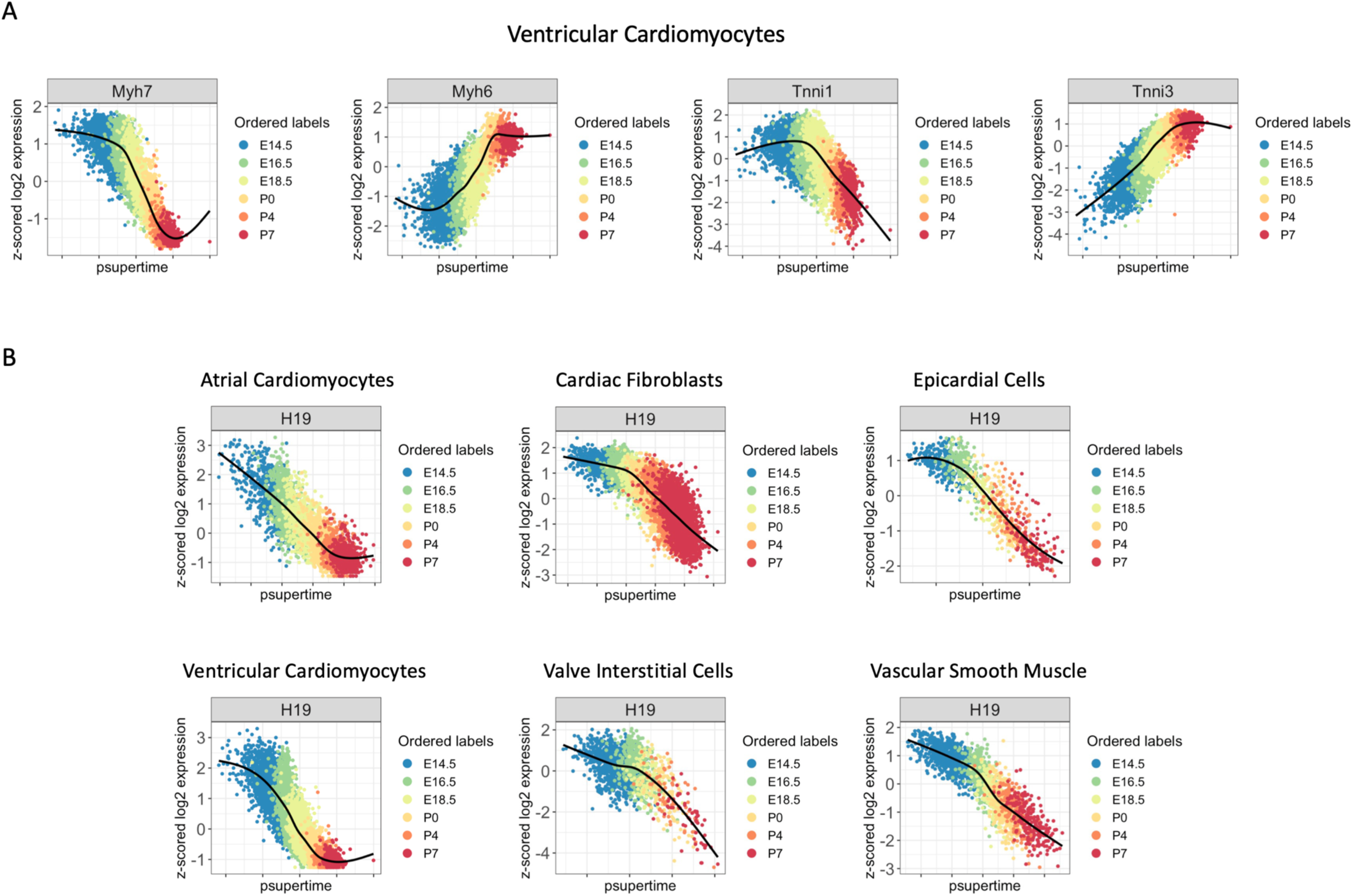
**A)** Plots of gene expression for *Myh6*, *Myh7*, *Tnni1* and *Tnni3* against pseudotime scores learned by psupertime in ventricular cardiomyocytes. x-axis = pseudotime score for each cell, y-axis = z-scored log2 gene expression values. All cells are ordered by pseudotime score and coloured by their known time point so as to easily visualise changes in gene expression across time. For example, a notably sharp drop in expression of *Myh7* can be seen from E18.5-P7, while *Myh6* steadily increases in expression from E16.5 to P4. This is consistent with known expression patterns. **B)** Plots as in Figure 6A of gene expression for *H19* against pseudotime scores learned by psupertime in multiple cardiac cell types. x-axis = pseudotime score for each cell, y-axis = z-scored log2 gene expression values. *H19* expression decreases consistently across time in cell types of distinct lineages including cardiomyocytes, epicardial cells and valve interstitial cells.

The number of dynamic genes per cell type varied significantly, ranging from 27 (LYMPHE) to 1556 (CFIB1). Fibroblasts and atrial/ventricular cardiomyocytes showed by far the most genes with dynamic expression over pseudotime, potentially reflecting a greater degree of perinatal development compared to other cell types. Fibroblasts are known to have extensive postnatal proliferation and involvement in ECM remodelling, while cardiomyocytes undergo significant changes previously discussed including hypertrophic cell growth with concomitant cell cycle arrest, sarcomeric protein isoform switching, and a transition to aerobic respiration (Padula et al., 2021; Uscategui Calderon et al., 2023). Caution must be taken with this interpretation however, as we noted positive correlation between cell number and the number of dynamic genes found per cell type (**Figure S4**). This is not unexpected as greater sample numbers generally allow for more statistically significant results.

On average we detected 404 genes with dynamic expression over pseudotime per cell type. Of the 4,886 genes identified in total, 3,737 (76%) were unique to one cell type and 234 (4.8%) were present in >= 3 cell types. The latter included mitochondrial genes, ribosomal genes, solute carrier (Slc) proteins, genes associated with metabolic pathways such as cyclooxygenase (Cox) and fatty acid binding protein (Fabp) family members, and some sarcomere genes. Low-level expression of *Mlc2a*, *Nkx2-5,* and *cTnt* has been previously observed in the pro-epicardial organ during development (Filipczyk et al., 2007; Mommersteeg et al., 2007) and could help explain the latter finding. The genes that most stood out overall in their prevalence across diverse cell types included *H19* (9 types), *mt.Nd1* (9 types), *Calm1* (8 types) and *Cd9* (7 types) (**Table S4**). Of those, the lncRNA H19 was the top psupertime hit (based on absolute beta coefficient scores) for epicardial cells, vascular endothelium cells, vascular smooth muscle cells, and macrophages, and in the top 5 for atrial cardiomyocytes, fibroblasts, and valve interstitial cells (**Figure 6b**).

### Decrease in lncRNA H19 expression is a common feature of cardiac cell maturation, with the exception of endocardial cells

Given the psupertime results, we further examined H19 expression patterns during cardiac cell maturation. H19 expression decreased strongly in most cell types, of distinct origins, from E14.5 to P7 (**Figure 7a**). Interestingly, endocardial and valve endothelial cells (known to be derived from endocardial cells) were the notable exception to this pattern, showing relative maintenance of H19 expression. To validate this finding, we also analysed a recently published, independent dataset (Feng et al., 2022) generated using a different single-cell technology called MULTI-seq. The same pattern was indeed replicated in this dataset, which included an expanded series of timepoints from E10.5 to P9. In most cases an initial increase in H19 expression peaking around E14.5 and decreasing thereafter could be seen. Consistent with our data, H19 expression was substantially maintained in endocardial cells and VECs (**Figure 7b**). We additionally analysed only male or female samples to verify that this pattern was not sex-dependent (**Figure S5**).

**Figure 7:**
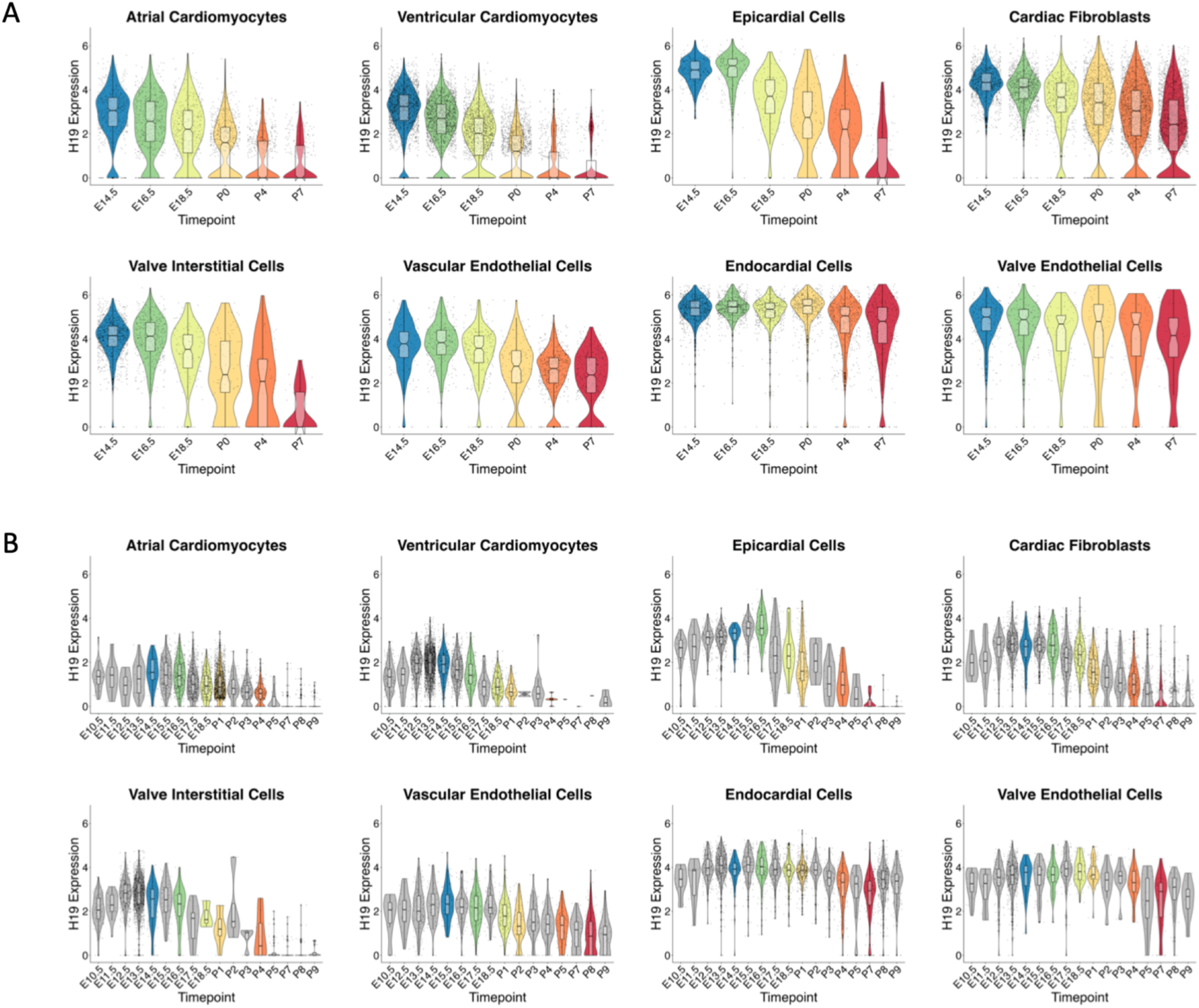
**A)** Seurat violin plots of *H19* gene expression per time point in multiple cardiac cell types. *H19* decreases in the majority of cell types, but appears to maintain expression in endocardial and valve endothelial cells. **B)** As above, but from the public dataset by Feng W. et al. spanning a wider developmental time period. *H19* initially increases early in development, peaks around E12.5-E14.5 and decreases thereafter in most cell types, but again is maintained in endocardial and valve endothelial cells.

To experimentally validate these *in-silico* results, we performed RNAscope staining using our earlier cell type specific markers alongside H19 **(Figure 8)**. H19 expression was generally diffuse but most highly expressed in endothelial cells, as expected from the scRNA-seq analysis. In each case a substantial decrease in H19 was seen over time in non-endocardial cells, with the greatest drop occurring from P1 to P7. Notably, by P7 H19 appeared to only remain in endocardial and valve endothelial cells, confirming maintenance of expression.

**Figure 8:**
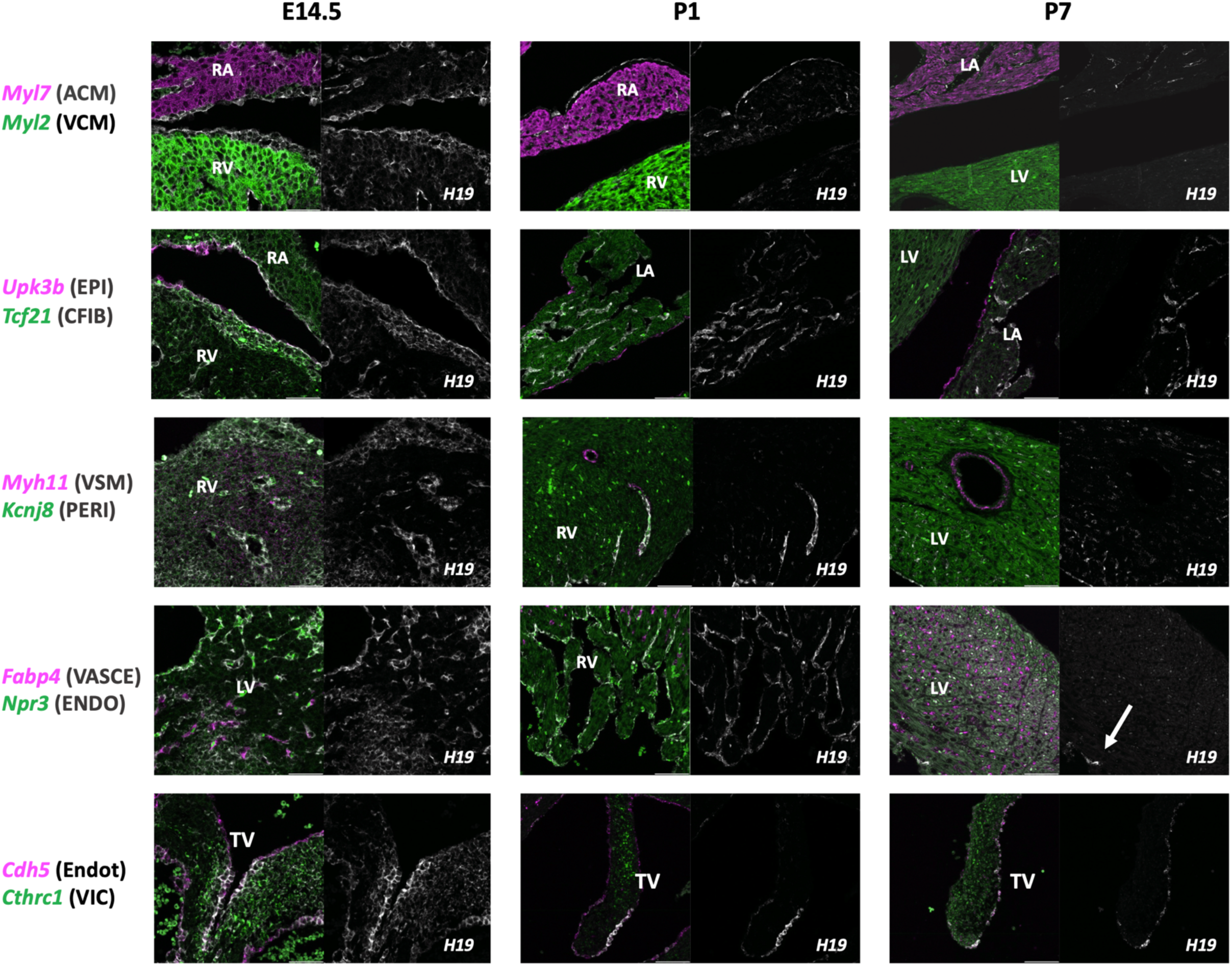
RNAscope stains across E14.5, P1 and P7 timepoints for *H19* and the same cell type marker genes as in Figure 3. At E14.5, H19 expression is relatively high and is co-expressed with all eight markers. A significant decrease in H19 expression thereafter is consistently seen, with the largest drop from P1 to P7. Note that the remaining expression of *H19* at P7 corresponds to endocardial and valve endothelial cells, consistent with apparent maintenance seen in our scRNA-seq data. RA = right atrium, RV = right ventricle, LA = left atrium, LV = left ventricle, TV = tricuspid valve.

Interestingly, the RNAscope analyses showed valve H19 expression becoming more spatially localised over time. At E14.5, H19 was diffusively expressed in VICs and present in VECs on both sides of the valve. However, at P1 and P7, H19 appeared to be specifically expressed in VECs and only at the coapting interface. This highlights the importance of supplementing scRNA-seq data with staining when analysing gene expression in order to obtain precise spatial information.

### Shared expression dynamics highlight imprinting as an important regulator of perinatal growth

Having identified H19 as a potential regulator of maturation in the heart, we wanted to investigate potential downstream effects of H19 and other genes showing dynamic expression during the perinatal period. We thus performed enrichment analysis of 1,149 genes identified by psupertime as showing significant dynamic expression in at least 2 different cell types, using the MGI Mammalian Phenotype 2021 library. This analysis revealed significant (Benjamini-Hochberg adjusted p-value <0.05) enrichment for 37 terms (**Table S5**). The top three terms in order of descending p-value were cardiac hypertrophy (MP:0001625), increased heart weight (MP:0002833), and dilated heart left ventricle (MP:0002753), with many of the remaining significant terms also referencing dilation or hypertrophy. These results are consistent with the cell growth-related TFs and pathways found in the decoupleR analysis. While these terms reference pathological processes, many of the genes driving the results are necessary for normal heart development. One example from our analysis is *Cav1*, which is expressed primarily in fibroblasts and endothelial cells and is required for normal formation of vesicular invaginations of the plasma membrane called caveolae (Liu et al., 2008). *Cav1* null mouse hearts demonstrate significant cardiac hypertrophy, likely due to hyperactivation of the p42/44 MAP kinase cascade which *Cav1* negatively regulates (Cohen et al., 2003). A number of cardiac developmental genes are also in fact known to cause hypertrophic pathology when improperly reactivated (referred to as the “foetal gene program”) (Kuwahara et al., 2012; Olson and Schneider, 2003).

A search for H19 within the Enrichr results found it to be associated with decreased body weight (MP:0001262) (p=0.013). It was also associated with increased placenta weight (MP:0004920), paternal imprinting (MP:0003123) and genetic imprinting (MP:0003121), all of which had high-ranking enrichment scores (combined p-value/odds ratio). The latter terms reflect that H19 is known for being part of the major imprinted H19/IGF2 locus, with *H19* being expressed exclusively from the maternal allele and *IGF2* from the paternal allele. Maternal imprinting (MP:0003122) was itself significant (p=0.03) in the enrichment analysis. We therefore reviewed the psupertime expression patterns of imprinted genes from both sets and found that VECs also showed remarkably less change in expression in many of these genes compared to other cell types such as epicardial cells, fibroblasts, and ventricular cardiomyocytes (**Figure 9**). This suggests that maintenance of imprinted genes may be of particular importance for VEC maturation.

**Figure 9:**
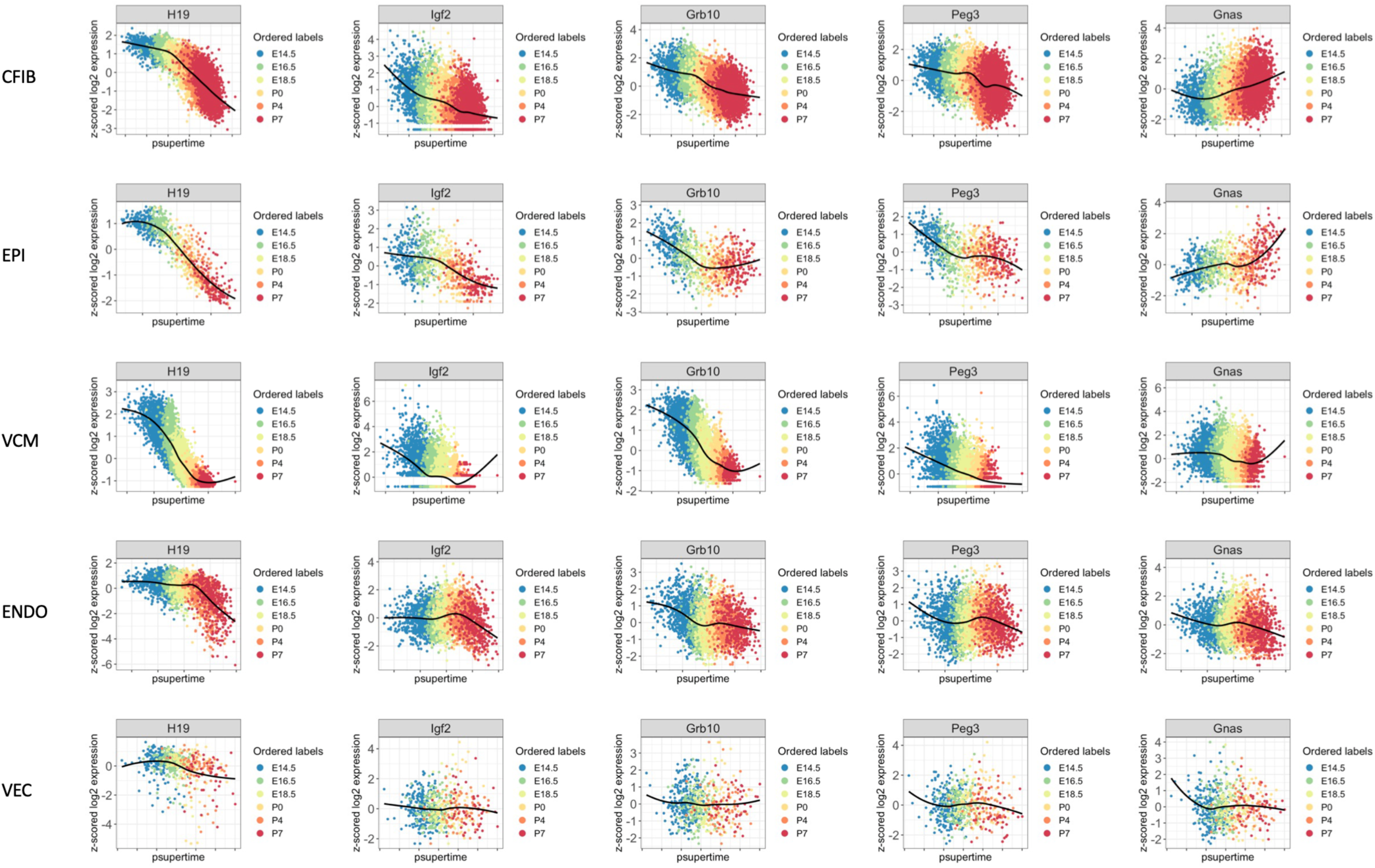
Plots as in Figure 6 of gene expression for *H19, Igf2, Grb10, Peg3 and Gnas* against pseudotime scores learned by psupertime in multiple cardiac cell types. x-axis = pseudotime score for each cell, y-axis = z-scored log2 gene expression values. These genes were identified from the imprinted gene sets found to be highly enriched over maturation (**Table S5**). In most cell types a notable change in expression of these genes can be seen, often decreasing over time. However by comparison, less variation in expression is seen in endocardial and especially valve endothelial cells.

## Discussion

In this study we performed high-throughput, droplet based single-cell transcriptomics of mouse hearts at six different time points throughout the perinatal period (E14.5 to P7). We identified major cell types such as cardiac fibroblasts, cardiomyocytes, and endothelial cells, but were also able to sample much less frequent cells, such as melanocytes, neuronal cells, and lymphatic endothelial cells. Our dataset vastly increases the number of cell transcriptomes and cell types characterised compared to previous studies that included foetal to neonatal transitions. This largely increased sample size allowed us to determine the common transcriptional response of cardiac cells during the transition to neonatal life. We found that the great majority of enriched GRNs and pathways were related to regulation of cell growth and proliferation. Notably Fos/Jun, STAT3, KLF, and NF-Kβ (upregulated in maturation) are all known to directly modulate transcription of Cyclin D1 (Duronio and Xiong, 2013), a nuclear protein required for cell cycle progression in G1. Cyclin D1, along with other positive cell cycle regulators such as Rb, is known to be highly expressed in foetal cardiomyocytes but significantly downregulated in neonatal cardiomyocytes, coinciding with postnatal cell cycle arrest (Zhao et al., 2020). c-MYC (downregulated with maturation) forms a heterodimeric complex with the protein MAX which activates a variety of known target genes essential for cell-cycle progression and growth including Cyclin D2, CDK2, and the translation initiation factors eIF4 and eIF2 (Pelengaris et al., 2002).

We observed a decrease in Skil activity over time across many cardiac cell types corresponding with increased Smad2/3 and resulting TGFβ pathway activity. The TGFβ pathway is highly important for development and multifunctional, with context-dependent effects on cell growth (Massagué, 2012). A prior study found that a small set of master transcription factors direct Smad3 to cell type-specific binding sites and determine cell type-specific responses to TGFβ signalling (Mullen et al., 2011). Similar cases for other pathways may help explain why our results appeared less cell-specific than perhaps anticipated, given the number of distinct cardiac cell types that must be specified in development. Given the known association of TGFβ with heart disease (Frangogiannis, 2022), it is of high interest to further study the impact of increased TGFβ activity on each cell type throughout cardiac maturation.

It is worth stressing that the Hallmark pathways identified by decoupleR do not act in isolation, and can in fact regulate each other. For example, the IL-6-STAT3 pathway is known to mediate the prevention of apoptosis induced by TGFβ (Hirano et al., 2000). Interestingly, both STAT3 and NF-Kβ target and drive expression of c-Myc (Duyao et al., 1990; Kiuchi et al., 1999) and may thus be necessary to prevent complete downregulation of c-Myc during maturation.

Analysis of time-varying genes via psupertime likewise showed a common enrichment in terms relating to cell growth. Given the relative commonality seen in the pathway results, it is curious that most of the dynamic genes were actually cell-specific. It must be kept in mind though that high-level pathways (as used by decoupleR and prevalent in development) are themselves rarely specific and typically associated with a large number of genes, thus increasing statistical significance. The overall top time-varying gene was the lncRNA H19, part of a major imprinting locus with IGF2. H19 significantly decreased across multiple cell types in our analysis as well as in an independent dataset, and this was further verified experimentally via RNAscope.

The identification of a signature for imprinted genes suggests that common gene expression changes across cell types in the murine heart during perinatal development are involved in regulation of cell and organ growth after birth. While the downregulation of imprinted genes across different organs postnatally has been shown previously (Lui et al., 2008), to the best of our knowledge this is the first observation of this mechanism across different cell types in the mouse heart during development. The reason for the downregulation of imprinted genes such as H19 in many organs postnatally is still strongly debated (Moore et al., 2015; Moore and Haig, 1991). Our findings indicate that downregulation of imprinted genes can be observed across multiple cell types in the newborn heart and coincides with transition from foetal to neonatal growth in the heart.

The H19/IGF2 locus has been well studied and known to be involved in foetal programming through DNA methylation. In brief, the locus is regulated by an imprinting control region (ICR) containing binding sites for the insulator protein CTCF (Szabó et al., 2004). Unmethylated ICR on the maternal allele allows binding of CTCF, which blocks transcription of IGF2 and activates H19, whereas on the paternal allele, ICR methylation activates IGF2 transcription and silences H19. Misregulation of this locus causes growth disorders in humans including Beckwith-Wiedemann syndrome and Silver-Russell Syndrome (Bliek et al., 2006; Brunkow and Tilghman, 1991; Gabory et al., 2010; Sparago et al., 2004). These conditions, particularly the former, are known to be associated with cardiovascular defects including congenital heart disease (Ghanim et al., 2013; Greenwood et al., 1977). In general, IGF2 has a growth promoting effect while H19 has a growth restraining effect. H19 was in fact highlighted as anti-hypertrophic by a study in which ectopic induction of H19 expression was sufficient to suppress cardiomyocyte hypertrophy in vitro (Viereck et al., 2020). In recent years one study has positively linked cardiovascular pathology in mouse loss of imprinting models to misexpression of H19 and specifically reduction of H19 binding to Mirlet7 (Park et al., 2021). However, the molecular disease processes of the above growth syndromes have not yet been extensively studied or examined in the context of perinatal heart maturation.

In our study, endocardial cells appeared to be an exception to the observed overall downregulation of H19, instead showing relative maintenance of H19 expression. The majority of heart valve development takes place between E9.5 and E14.5 (Feulner et al., 2022) and previous reports suggested that subsequent endocardial contributions are restricted by a VEC-specific Nfatc1 enhancer (Zhou et al., 2005). Nfatc1 modulates endothelial-to-mesenchymal transition (EndoMT) in a cell-autonomous manner by suppressing transcription of Snail1/2 (Wu et al., 2011). This is relevant as interaction of H19 with the polycomb repressive complex 2 suppresses H3K27 tri-methylation of the Tescalcin locus, leading to reduced NFAT expression and activity (Viereck et al., 2020). H19 may thus play a role in limiting Nfatc1 signalling by the autoregulated enhancer in VECs, ensuring normal balance between EndoMT and valve elongation. H19 also helps to prevent EndoMT via TGFβ (Thomas et al., 2019) and has specifically been found to inhibit TGFβ2-induced EMT of the eye lens by suppressing Smad-dependent signalling (Xiong et al., 2022). We therefore propose that stable H19 expression in VECs is required for maintenance of the VEC phenotype, and that misregulation of H19 might thus lead to reduced valve endothelium. Further experimental validation is required to determine whether H19 and Nfatc1-mediated regulation of VEC phenotypes and foetal valve maturation are related. In contrast, downregulation of H19 in other cell types is likely required for the continuous contributions of these cell types to early postnatal cardiac growth. Additionally, H19 overexpression in VICs has been found to alter the Notch1 pathway, leading to increased expression of Runx2 and development of an osteoblastic VIC phenotype resulting in calcific aortic valve disease (CAVD) (Hadji et al., 2016). The downregulation of H19 seen in VICs is thus likely important in maintaining their quiescence and preventing improper activation.

The results from our scRNA-seq dataset illustrate the complex interplay of GRNs and pathways necessary to balance proliferation (via modulation of cell cycle progression vs apoptosis) and hypertrophic cell growth during cardiac cell maturation. Multifaceted pathways like TGFβ that are capable of modulating all these functions in various contexts are thus understandably key. Our study additionally suggests that modulation of imprinted genes, in particular the lncRNA *H19*, is important for cardiac cell maturation. The overall downregulation of *H19* that we observed likely plays a hitherto-unappreciated role in maintaining normal proliferative levels and phenotype of many cell types including valve interstitial cells, with disease implication in cases of dysregulation. Conversely, our results suggest that stable *H19* expression is specifically required for phenotype maintenance of endocardial and especially valve endothelial cells.

### Limitations

Our study validated known cardiac cell types and markers, and we additionally identified novel candidate genes and pathways involved in the transition from foetal to neonatal life. However, our analyses (and those in other scRNAseq studies that require tissue dissociation) may be affected by capturing cardiomyocytes at a lower rate with increasing age, while less fragile cells, such as stromal and interstitial cells are relatively enriched. Consequently, maturing binucleated or multinucleated cardiomyocytes may be underrepresented. Another factor hindering capture of maturing cardiomyocytes is their increasing size, which can make them too big to be captured in droplet based single-cell applications.

## Materials and Methods

### Animal ethics and collection of heart tissue

All experiments involving mice were performed according to the Canadian Council on Animal Care (CCAC) guidelines and were approved by the ethical committee of the CHU Sainte-Justine Research Centre. Mice acquired from Jackson Laboratory (C57Bl/6NJ, stock nr.:005304) were maintained on a C57Bl/6NJ background at the CHU Sainte-Justine animal facility. Mice were kept on a 12 hour light/dark cycle (6.a.m to 6p.m.) under specific-pathogen-free (SPF) conditions, with ad libitum access to water and food on a standard chow diet (2918 Teklad global 18% protein rodent diets (Supplier Envigo)). Room temperature was maintained between 20-24℃ with 40-70% humidity, according to CCAC guidelines. Mice breedings were set up between wild-type C57Bl/6NJ mice. Matings were checked in the morning and if a copulation plug was present, noon of that day was counted as E0.5 of the pregnancy.

Methods for extracting foetal hearts were the same as in Wünnemann et al. (Wünnemann et al., 2020). Briefly, pregnant females were sacrificed by injection of Pentobarbital followed by cervical dislocation of the female. The uterus containing foetuses was extracted and kept on ice in DPBS during dissection. Fine dissection of individual foetuses was performed in cold DPBS under a stereoscopic microscope. For postnatal time points, the morning of birth was considered as P0 and pups were processed the same day, while P4 and P7 were processed four days and seven days following the morning of birth respectively. For all time points, the heart was extracted by cutting the abdominal aorta using scissors and carefully pulling out the lungs together with the connected heart. Any remaining lung tissue, trachea, fat or non-cardiac tissue was removed. The pulmonary veins and the inferior vena cava were cut as close to the base of the heart as possible and the pulmonary artery and aorta were cut close to the pulmonary valve and aortic valve, respectively, to exclude the majority of these two large vessels from the tissue dissociation.

### Single cell dissociation of heart tissue

Dissected hearts were cut into small pieces using fine dissecting scissors and transferred to gentleMACS C-tubes (Miltenyi Biotec, Cat-Nr:130-093-237) containing enzymatic digestion solution according to manufacturer’s protocol for the Neonatal Heart Dissociation Kit (Miltenyi Biotec, Cat-Nr: 130-098-373). Two mouse hearts were processed in separate C-tubes at the same time for all experiments presented in this manuscript. C-tubes were subsequently fitted onto a gentleMACS Octo Dissociator (Miltenyi Biotec, Cat-Nr: 130-095-937) using the program *37C_mr_NHDK_1*, which performs enzymatic and gentle mechanical tissue dissociation at 37°C according to manufacturer’s protocol. Following completion of the dissociation program, C-tubes were briefly centrifuged at 400g to collect all liquid and cells from the lid and walls of the tube. Cells were then treated according to the manufacturer’s protocol. Briefly, 7.5ml of DMEM High Glucose medium without phenol red (Thermo Fisher, Cat-Nr.: 21063029) containing 10% foetal bovine serum (FBS) preheated to 37℃ was added to the cells followed by resuspension by mild pipetting (5-10 times using a 10ml pipette). Cells in DMEM+FBS were subsequently filtered through a 70µM filter (Miltenyi Biotec, Cat-Nr: 130-098-458) and transferred to 15ml conical Falcon tubes. Filters were washed with 3ml warm DMEM+FBS.

Following filtration, cells were pelleted for 5 min at 600ɡ at room temperature. Supernatant was removed and cell pellets were resuspended in 1 ml PEB buffer (DPBS pH 7.4, 0.5% bovine serum albumin (BSA), 2mM EDTA) followed by addition of 10ml 1x red blood cell lysis (RBC) buffer (Miltenyi Biotec, Cat-Nr.: 130-094-183). Cells were incubated in RBC lysis buffer for 2 min at room temperature and then centrifuged at 600ɡ for 5 min at room temperature. Supernatant was removed and cell pellets resuspended in 10ml DPBS + 15µl of Enzyme A of the dissociation kit. Cells were then centrifuged for a last time at 600g and resulting pellets were resuspended in 1ml of DPBS + 0.01% BSA for counting. Cells were counted using a disposable Fuchs-Rosenthal hemocytometer (INCYTO C-Chip Disposable Hemocytometer) with trypan blue to assess viability. Following counting, cells were diluted to a final concentration of 140 to 160 cells/µl and processed using Drop-seq.

### Single-cell RNA-seq of mouse hearts using Drop-seq

Dissociated cells were captured in nanoliter droplets using a custom Drop-seq setup at the CHU Sainte-Justine Research Center in the Andelfinger laboratory. The custom Drop-seq setup and protocol closely followed the originally published protocol, with a few minor differences (https://mccarrolllab.org/dropseq/) (Macosko et al., 2015). We utilised Smith’s Medical Medfusion 2500 pumps in place of KD Scientific Legato 100 pumps as described in the original article. Cell dissociations were run through PDMS chips ordered with aquapel treatment from FlowJEM. To capture single cell transcriptomes, cells were run alongside droplet generation oil (Bio-Rad,Cat-Nr.: 186-4006 and barcoded beads (Chemgenes, CSO-2011) which were suspended in cell lysis buffer (see original Drop-seq protocol) supplemented with 50 µl of RNAse inhibitor (SuperASE, Cat-Nr.: AM2694) per ml of lysis buffer. The barcoded beads contain oligos with a bead-specific cell barcode, an oligo-specific unique molecular identifier (UMI) and a polyT sequence to capture polyadenylated RNA. Cells and beads were run at speeds of 4 ml/h, while oil was run at 15 ml/h. During each Drop-seq run, the aqueous flows from 1 ml of beads and cells were collected. Following droplet capture, droplets were diluted in in 30 ml 6x SSC (Thermo Fisher, Cat-Nr.: AM9763) and broken by shaking after adding 1ml of Perfluorooctanol (PFO, Fisher Scientific, Cat-Nr: AAB2015618). Beads were spun down and washed twice in 6x SSC and once in a home-made RT buffer. Beads were resuspended in RT mix (same reagents as original Drop-seq protocol) and subjected to reverse-transcription for 30 minutes at room temperature followed by 90 minutes at 42°C in a rotating hybridization oven. After reverse transcription, beads were washed, and treated with Exonuclease I for 45 minutes at 37°C with rotation. Following exonuclease treatment, beads were washed and counted using a Fuchs-Rosenthal hemocytometer in a 1:1 ratio with bead counting solution (10% PEG, 2.5M Nacl, as described in Gierahn et al 2017 (Gierahn et al., 2017).

To amplify single-cell transcriptomes, 4000 beads (∼200 single-cell transcriptomes attached to microparticles (STAMPS)) were used as input for individual PCR reactions as described in the original Drop-seq protocol. 10 PCR reactions were performed in parallel, for a total of 2000 STAMPs per biological sample. PCR reactions were pooled post-PCR for desired final number of STAMPS in the library and cleaned up twice using AMPure XP beads at 0.6x concentration. Quantity and size distribution of cleaned cDNA was checked using the Bioanalyzer High Sensitivity Assay (Agilent Genomics, Cat: 5067-4626). Libraries for sequencing were prepared from 600pg cDNA input using Nextera XT library preparation kit (Illumina, Cat: FC-131-1096) with i7 primers from Illumina (N70X) together with a custom P5 primer (see Drop-seq protocol). Quality control of libraries was performed using the Bioanalyzer High Sensitivity Assay. Libraries were quantified and quality controlled using the Universal Library Quantification Kit (KAPA) and were sequenced on an Illumina NExtseq 500 at the Institute for Research in Immunology and Cancer (IRIC) in Montreal. Read settings for sequencing were: R1 = 20bp, R2=63bp and index = 8bp.

### Processing and annotation of single-cell RNA sequencing data

Raw Illumina sequencing data was demultiplexed using bcl2fastq (v2.17). Fastq files for E14.5 are publicly available (GSE150817 (Wünnemann et al., 2020)) and were downloaded from the sequence read archive (SRA) using SRA explorer (https://sra-explorer.info/). Raw FASTQ files for each sample were processed using kallisto and bustools (kb-python, v0.24.4) against the ensemble reference release 98 for *Mus musculus* (GRCm38, primary assembly, see Colab notebook) (Melsted et al., 2021). We utilised emptyDrops from the DropletUtils package in R to call cells from the raw, unfiltered matrices from kb-python (FDR threshold <= 1%) (Lun et al., 2019). Genes expressed in less than 5 cells per sample were subsequently filtered out. To detect potential cell doublets, UMI matrices filtered based on emptyDrops were analysed per time point, using the scrublet package (v.0.2.3) in R via reticulate with default parameters and an expected doublet rate of 5% to calculate doublet scores. We subsequently removed cells with a scrublet doublet score over 0.25 as predicted doublets.

After sample level QC filters, sample matrices were merged into one joined Seurat object. We then applied general QC filters, removing cells with more than 10k UMIs and with less than 500 total genes detected. To exclude stressed and dying cells, we also excluded cells with more than 15% mitochondrial gene expression. After quality control steps, we ended up with a set of 68,912 cells that we used for a first round of clustering. We utilised sctransform (Seurat_4.0.0) via the *SCTransform* function in Seurat to normalise UMI counts for sequencing depth (Hafemeister and Satija, 2019). We included the percent of mitochondrial gene expression, cell cycle, and sex as variables to regress out in the sctransform normalisation and scaling step. For calculating cell cycle scores for regression, we translated human gene symbols from Nestorowa et al. (Nestorowa et al., 2016) to mouse gene symbols using the bioconductor biomaRt R package (Durinck et al., 2005) and subsequently assigned cell cycle scores and predicted cell cycle phase for all cells. We used S and G2M scores for each cell in the SCTransform step from Seurat to remove potential cell cycle effects from the dimensional reduction. After SCT normalisation, principal components (PC) were calculated in Seurat based on the 3000 most highly variable genes. We used 50 PCs for Uniform Manifold Approximation and Projection (UMAP) embedding of all cells into 2 dimensions (McInnes et al., 2020). Unsupervised cell clusters were identified using a shared-nearest neighbour (SNN) graph followed by the original Louvain clustering algorithm as implemented in Seurat FindNeighbors() and FindClusters() functions.

Potential marker genes in clusters assigned by Seurat were calculated using the wilcoxauc() function from the presto package (Ilya Korsunsky et al., 2019). After the first round of clustering, we removed additional cell clusters from the dataset. First, we noted one cluster of cells that mostly consisted of cells from 1 sample and expressed high levels of genes not generally expressed in the heart, but in the lung (marked by *Wfdc2* (L Bingle et al., 2020) and *Scgb3a2* (Naizhen et al., 2019)). These cells likely present contamination from non-cardiac tissue during dissection and were therefore removed from the final dataset. We further identified a cluster that was clustered with cardiomyocytes in high dimensional space but expressed a mix of many different cell type markers, showed low UMI content and high percent mitochondrial gene expression. We classified these to either be cell damaged cell multiplets or empty droplets with high ambient RNA content and therefore manually removed them. Finally, two clusters represented foetal and postnatal red blood cells (marked by gamma-globin (*Hbb-y*) and Haemoglobin subunit epsilon-Y2 (*Hbb-bs*), respectively) that remained after red blood cell lysis. Since red blood cells are not of interest in this study, we excluded them from the final dataset. 67,330 cells remained following the initial filter.

A second more stringent round of QC was subsequently performed using parameters specific to each timepoint and in some cases replicate, with the goal of eliminating remaining outliers. In particular, it could be seen that while a mitochondrial cutoff of 15 was often appropriate for postnatal cells, a lower cutoff was more optimal for earlier cells. Overall E14.5 cells were filtered for 500-2500 genes, 1000-5000 counts and percent mito 0.5-5; while P7 cells were filtered for 500-2000 genes, 1000-8000 counts and percent mito 0.5-8/15. This additional filtering resulted in a final dataset of 49,769 cells covering six time points (each with six replicates), from which we ran SCTransform normalisation again on raw UMI counts and re-calculated the clustering to compute final cell embeddings. Our final clustering was derived using RunUMAP with dims 1:83 and FindClusters at various resolutions ranging from 0.3-10. Stable marker genes for major cell types across the sampled time points were calculated using Seurat’s FindMarkers() function with min.pct = 0.25, min.dif.pct=0.2, only.pos=T, assay=“RNA”.

### Subsetting and reclustering of cell types

Each respective cell type was subsetted from the final Seurat dataset and converted to raw data using the DropletUtils package, which was then re-clustered to verify cell type purity and manually remove any outlying noncorresponding (ex. *Myl2* containing) clusters before continuing analysis. For each cell type, PCA DimPlots grouped by time point were examined and used to guide whether to merge subtypes for further downstream analysis. Because CFIB2 and VSM2 populations were primarily postnatal and appeared to form different trajectories on PCA plots (**Figure S6**), CFIB1 and VSM1 clusters alone were used for analysis of fibroblast and smooth muscle populations. ACM1 and ACM2 populations however appeared to form a more continuous trajectory. This could be due to fewer transcriptional differences between the two subtypes throughout maturation, and possibly indicating cell state changes, as opposed to the more substantial changes in fibroblasts or smooth muscle subtypes. Therefore, we merged ACM1 and ACM2 for further analysis. MAC1-3 were likewise merged. While there was some separation between the three macrophage populations, it was felt that this cell type was sufficiently distinct from the rest and thus combining them would not skew the downstream results.

### Pseudotime analysis of single-cell data

To identify genes that are relevant for the progression of cells from E14.5 to P7, we used ordinal logistic regression, as implemented in the psupertime package (https://github.com/wmacnair/psupertime) (Macnair et al., 2022) and followed a similar approach as presented by Cook and Vanderhyden (Cook and Vanderhyden, 2020). We first estimated pseudotime using standard log-normalised UMI counts (RNA@data slot) from the Seurat object as input for psupertime. Pseudotime was calculated separately for each cell type that had at least 10 cells per time point sampled and cells identified in at least 5 out of 6 time points. This filter excluded neuronal cells, B-cells and T-cells from the time series analysis. Psupertime was run with penalization = “best”, testing all genes identified by Seurat in the given cell-type. The result was an R psupertime object in which the genes showing significant dynamic change in expression over time were contained (specifically, psuper_object$beta_dt$symbol). These genes could be shown using the function plot_identified_genes_over_psupertime.

Enrichment testing for a common maturation signature was performed on genes that were significantly differentially expressed in more than 2 different cell types against the Enrichr (Chen et al., 2013; Kuleshov et al., 2016; Xie et al., 2021) MGI Mammalian Phenotype Level 4 2021 library at https://maayanlab.cloud/Enrichr.

### decoupleR transcription factor and pathway activity analysis

The tool decoupleR was used to infer transcription factor activity across time for each cell type using CollectTRI, a curated database of TFs and their downstream targets. We performed a univariate linear model analysis using the run_ulm function, with the “mat” parameter being the log-normalized data matrix and the “net” (network) parameter being the CollectTRI database. This fit a linear model for each cell in our dataset and each TF in our network that predicted the observed gene expression based solely on the TF’s TF-gene interaction weights. The obtained t-value of the slope after fitting was the activity score. Positive scores indicated the TF was active while negative scores indicated the TF was inactive and 0 indicated lack of regulation by the TF. These scores were then imported into the corresponding seurat object as a separate assay and scaled. The mean scores for each timepoint per cell type were calculated and standard deviation was used to assess TF activity variation across time. The mean activity scores per time point of the TFs with the greatest standard deviation (ie. showing the most variation in activity) were then plotted using Complex Heatmap.

A similar process was used for pathway analysis. In this case, an overrepresentation analysis was performed using the run_ora function, with “mat” being the log-normalized data matrix and “net” being the Hallmark dataset from msigdb. For each cell and Hallmark pathway, an overrepresentation score was obtained corresponding to the -log10 p-value from a one-tailed Fisher’s exact test. These scores were added to the Seurat object for the cell type as a new assay and scaled. As with the TF analysis, the mean scores for each timepoint per cell type were calculated and standard deviation was used to assess pathway activity variation across time. The mean overrepresentation scores per time point of the top resulting pathways (showing greatest variation across time) were plotted using Complex Heatmap. An additional analysis using the PROGENy database was likewise performed, in this case using a multivariate linear model via the run_mlm function.

### SCENIC gene regulatory network analysis

Gene regulatory networks were constructed using the SCENICprotocol nextflow pipeline (https://github.com/aertslab/SCENICprotocol, Pipeline version: 0.2.0, pyscenic_container = aertslab/pyscenic:0.9.19) (Aibar et al., 2017; Van de Sande et al., 2020). The corrected UMI counts from the seurat object were converted to a loom object using the Convert() function from the loomR package and used as input for SCENIC using three cisTarget databases of TF binding motifs for scoring regulons (mm9-tss-centered-10kb-7species.mc9nr, mm9-tss-centered-5kb-7species.mc9nr, and mm9-500bp-upstream-7species.mc9nr). Regulon specificity scores (RRS) were calculated using the regulon_specificity_scores function in pySCENIC, which is based on Jensen-Shannon Divergence and described in Suo et al 2018 (Suo et al., 2018). The RSS measures how close the Regulon AUC score is to the case where it is only active in one cell type, ie. fully specific.

### Data and statistical analysis

Data analysis and statistical tests in this work were performed using R (<=4.2.3) and Rstudio (<=2023.06.0+4.1) (©Rstudio Inc).

### RNAscope

Preparation of samples and pre-treatment were performed according to the instructions in the RNAscope Multiplex Fluorescent Reagent Kit v2 user manual (https://acdbio.com/documents/product-documents). C57BL/6 hearts were collected and fixed overnight for paraffin embedding. Paraffin sections of 5μm thickness were heated for 1 h at 60 °C. Paraffin was removed from the slides by two Histo-Clear washes for 5 min and were dehydrated twice in 100% ethanol for 5 mins. Slides were incubated with hydrogen peroxide for 10 min at room temperature and a manual target retrieval (Appendix B of the protocol) was performed for 15 min at 95°C and then slides were left to dry. RNAscope Protease Plus was added for 30 min and incubated at 40 °C in the HybEZ Humidifying System Oven (Advanced Cell Diagnostics). Slides were washed twice for 2 min with distilled H20. The probes were heated to 37 °C for 30 min and cooled down to RT. One volume of C2, C3, and/or C4 probes were added to 50 volumes of C1 probe or TSA Buffer. The probe mixes were incubated for 2 h at 40 °C and the slides were washed twice for 2 min with Wash Buffer. All probes used were manufactured by Advanced Cell Diagnostics for the Manual Assay RNAscope and included: Adamts19 (500981; C4), Cdh5 (312531; C1), Cthrc1 (413341; C4), Fabp4 (884621; C2), H19 (423751; C3), Hapln1 (448201; C2), Kcnj8 (411391; C1), Myh11 (316101; C2), Myl2 (527951; C2), Myl7 (584271; C1), Npr3 (502991; C2), Tcf21 (508661; C1), Upk3b (568561; C1). All ensuing incubations were made at 40°C in the HybEZ oven; RNAscope wash buffer was used between each step. Incubation for AMP1 and AMP2 was 30 min, followed by AMP3 for 15 min. The horseradish peroxidase (HRP) was developed by using the RNAscope Multiplex FL V2 HRP-C1, -C2, -C3, and/ or -C4 reagent for 15 min followed by incubation with an opal dye (Opal 570; FP1487001KT; Perkin Elmer, Opal 620; FP1488001KT, Perkin Elmer, or Opal 690; FP1497001KT; Perkin Elmer). The HRP signal was blocked by adding the RNAscope Multiplex FL V2 blocker for 15 min. After washing, slides were counterstained with DAPI for 1 min at RT and mounted with Fluorescence Mounting Medium (S3023; Dako). Fluorescence was observed with a SP8-DLS confocal microscope (Leica Microsystems). Images were taken with the 20X objective and 2X zoom (final 40X magnification), total Z-size of 12μm (number of steps: 7; Z-step size: 2μm) and were tiled with 20% overlap. Image merging and projection was done with LASX imaging software (Leica, version 3.3.0.16799).

## Data and code availability

Raw FASTQ files and a processed, joined data matrices of all final cells used in the manuscript are to be available from NCBI Geo (GSE150817). Raw data for E14.5 hearts has been published previously (GSE109247) (Wünnemann et al., 2020). We selected six wild-type replicates (GSM2935824, GSM2935825, GSM2935826, GSM2935827, GSM3677518, GSM3677519) from the E14.5 dataset for equal number of replicates across all time points. Our data will be uploaded to Chan Zuckerberg CELLxGENE Discover (https://cellxgene.cziscience.com) for visual exploration. Code scripts and R files will additionally be available in future at https://github.com/larafeulner.

## Acknowledgements

This research was enabled in part by support provided by Calcul Quebec (https://www.calculquebec.ca/en/) and the Digital Research Alliance of Canada (https://alliancecan.ca/en). We would like to thank Melissa Goldman from the McCarroll Lab for the Drop-seq demonstration and home-made buffer RT recipe, J. Huber at the IRIC genomic platform for performing the Illumina sequencing for the Drop-seq libraries, and P. Gendron at the IRIC Bioinformatics platform for data demultiplexing. We also want to thank S. L’Espérance, K. Jolibois-Ouellete and D. Deraspé for their help and support with the mouse work. We would like to thank Michel Barrette with IT infrastructure. Finally we thank the Plateforme d’Imagerie Microscopique at CHU Sainte-Justine, including Elke Küster-Schöck, for their assistance with the RNAscope experiments.

## Supplementary Figures

**Figure S1:**
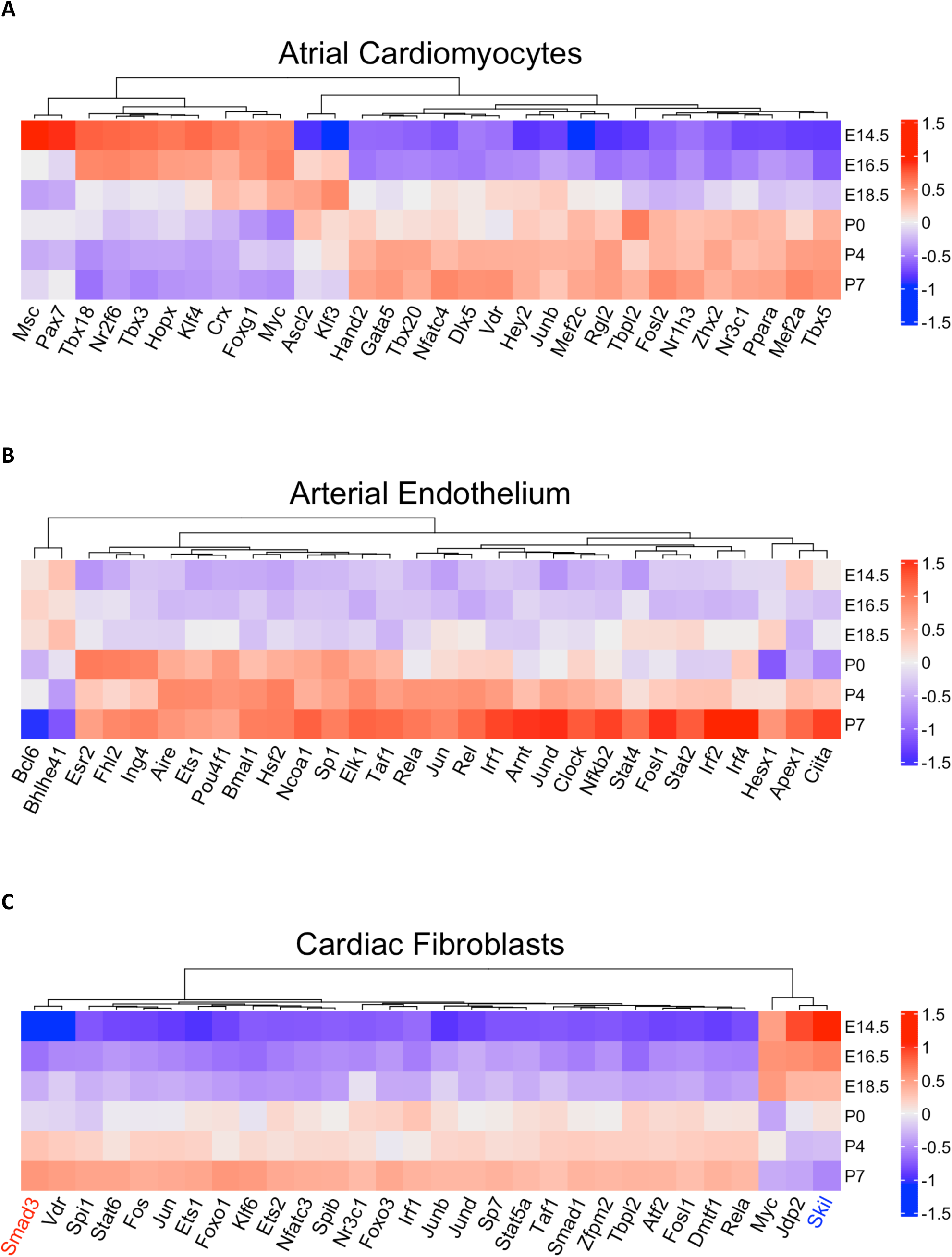
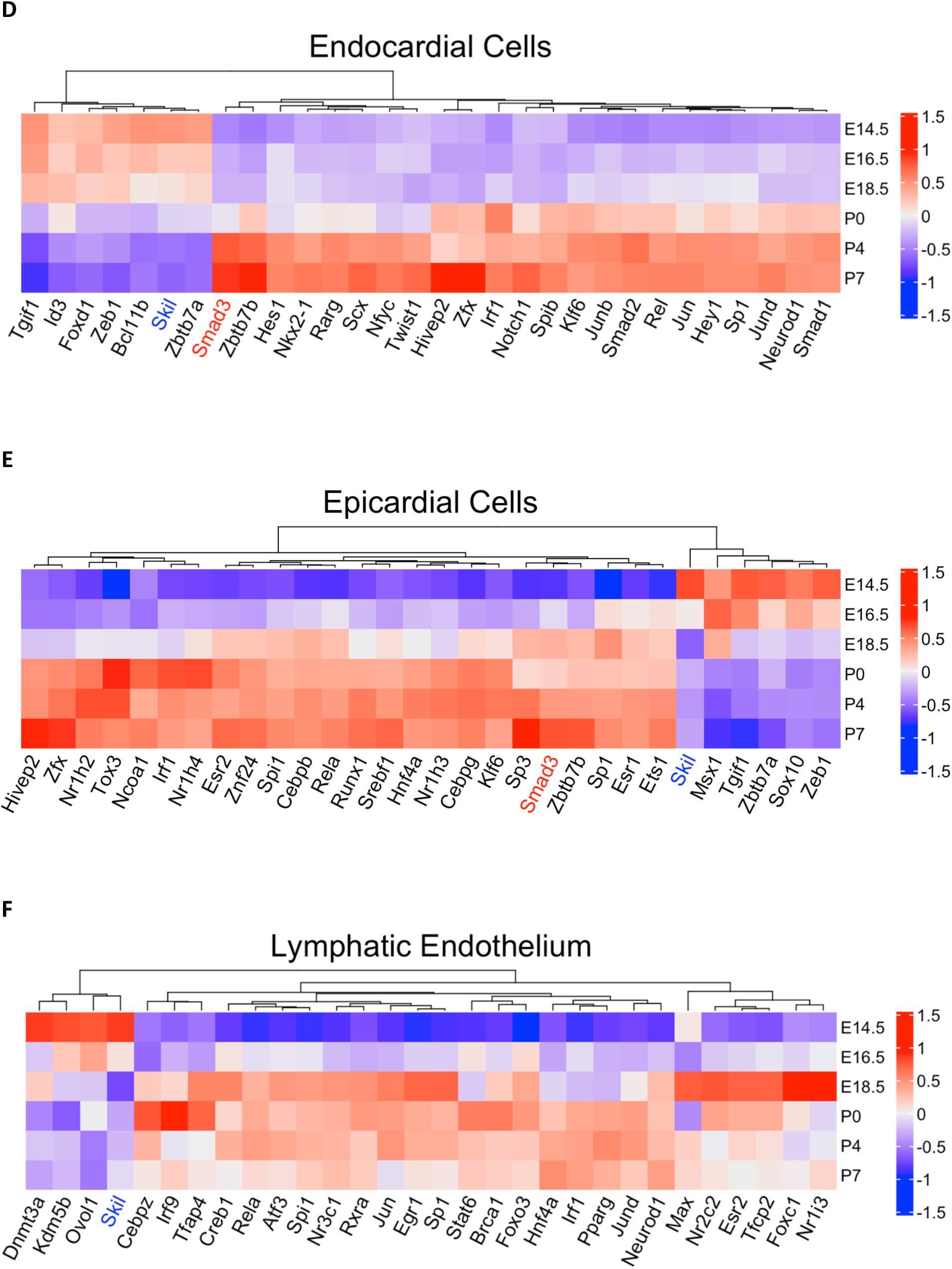
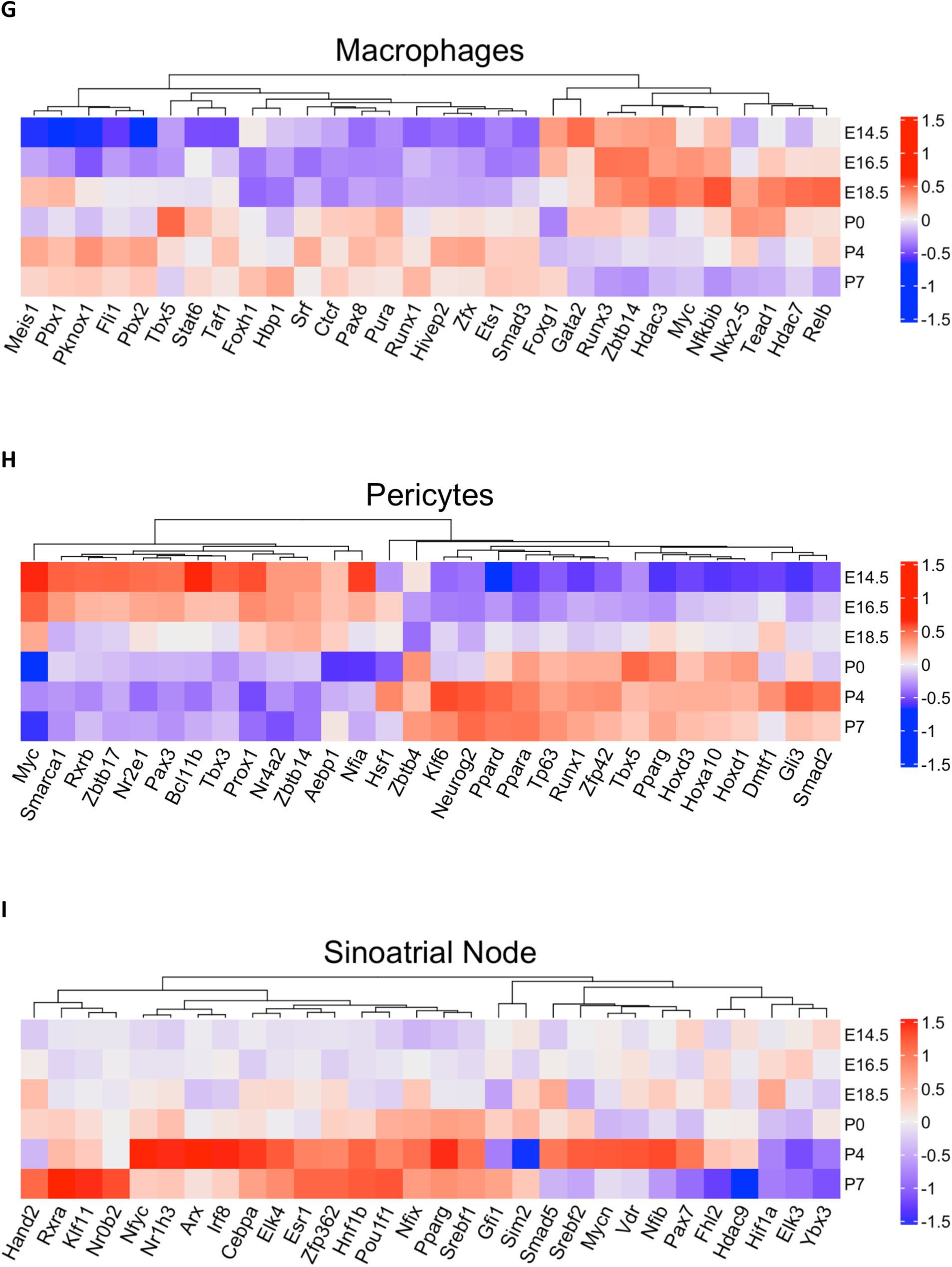
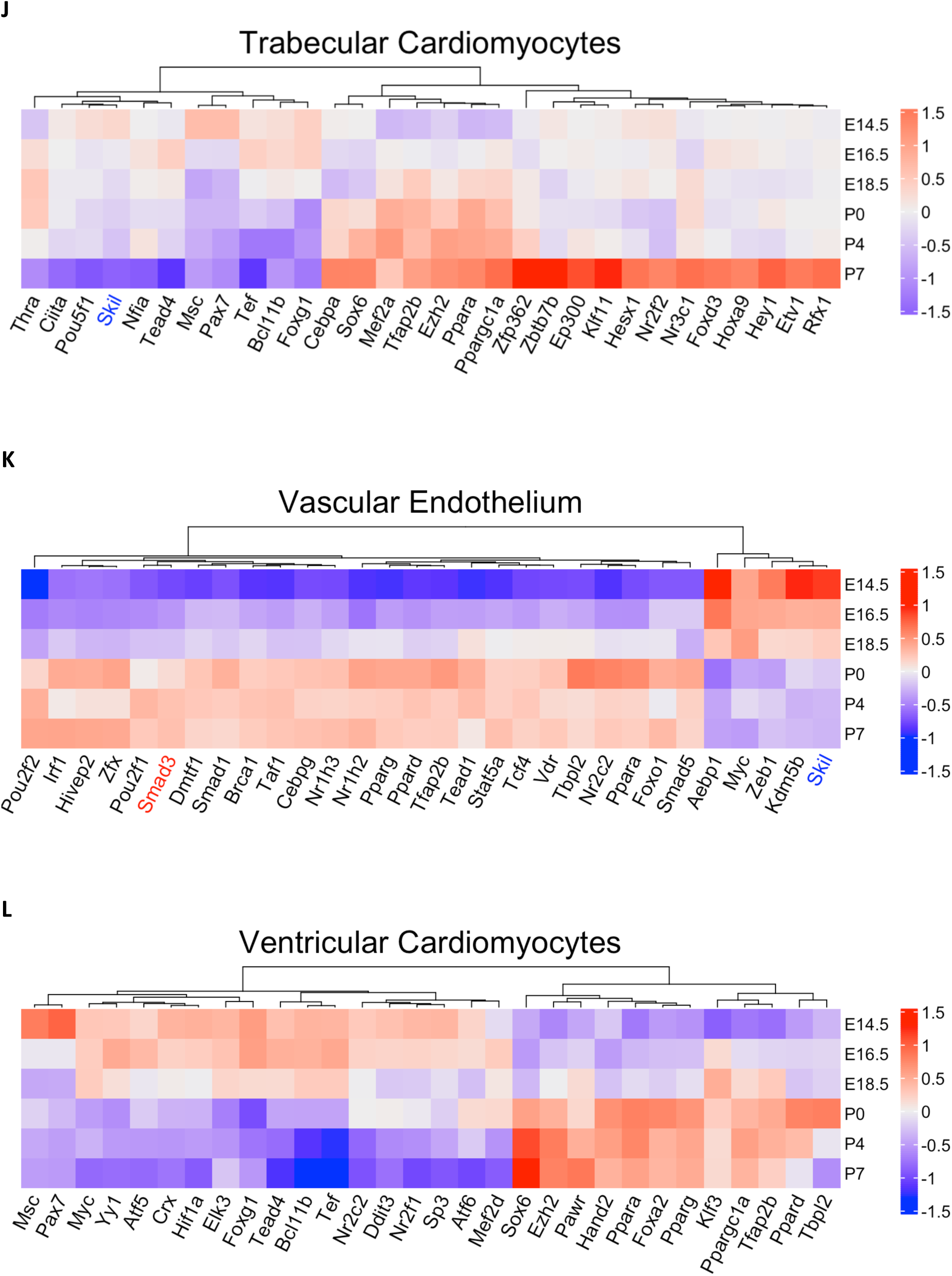
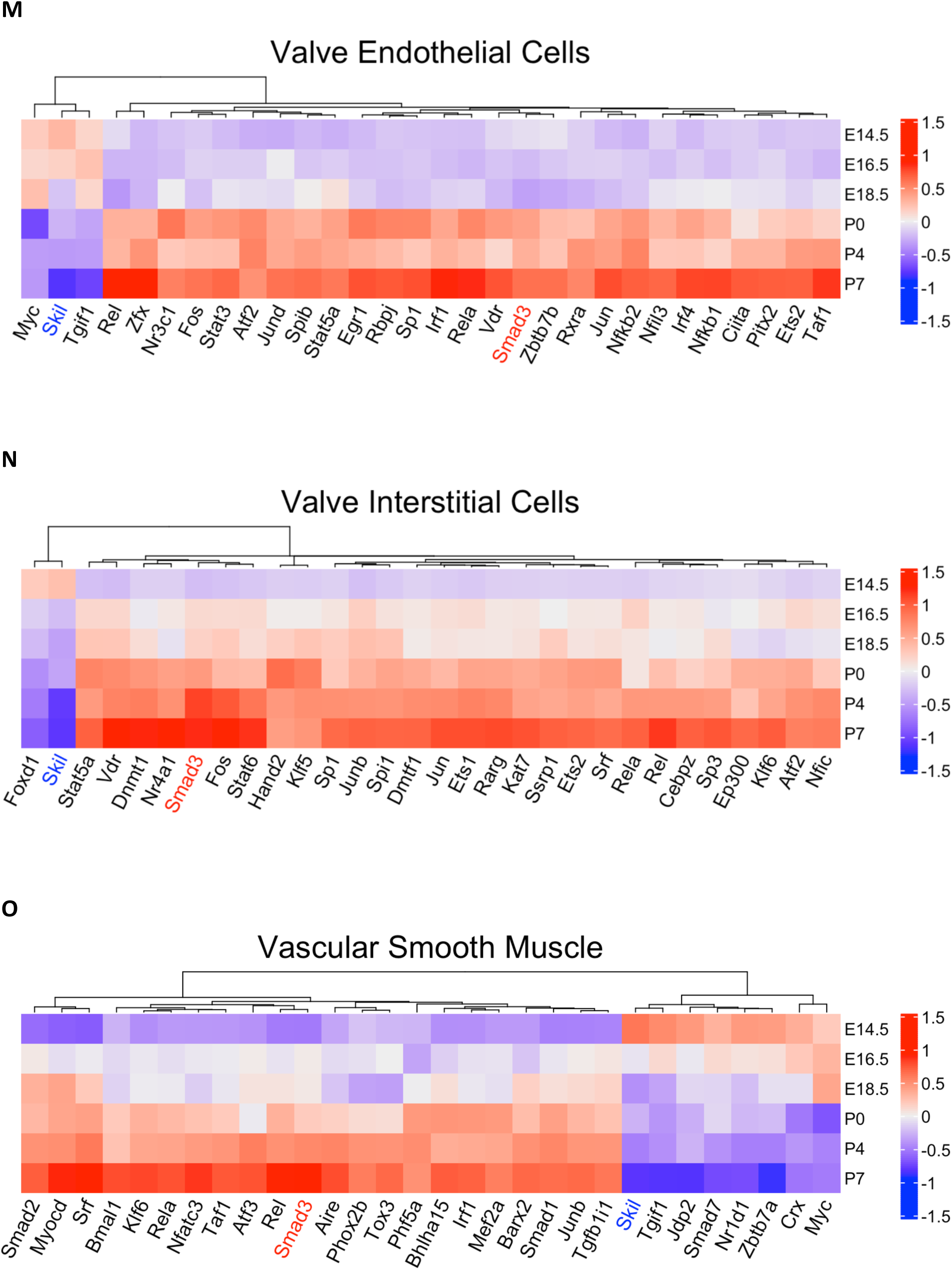
decoupleR TF activity heatmaps as in Figure 4a for all analysed cell types.

**Figure S2:**
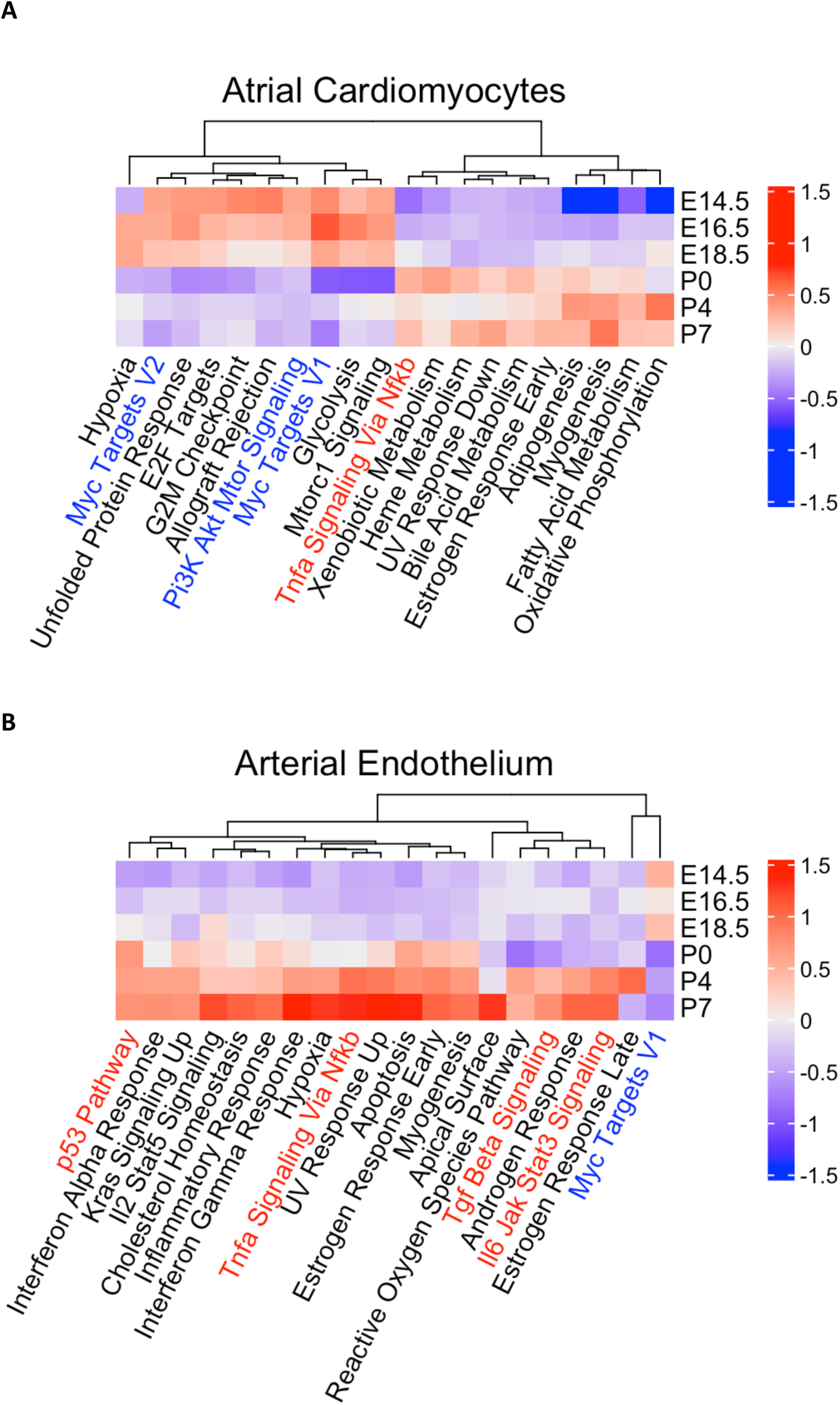
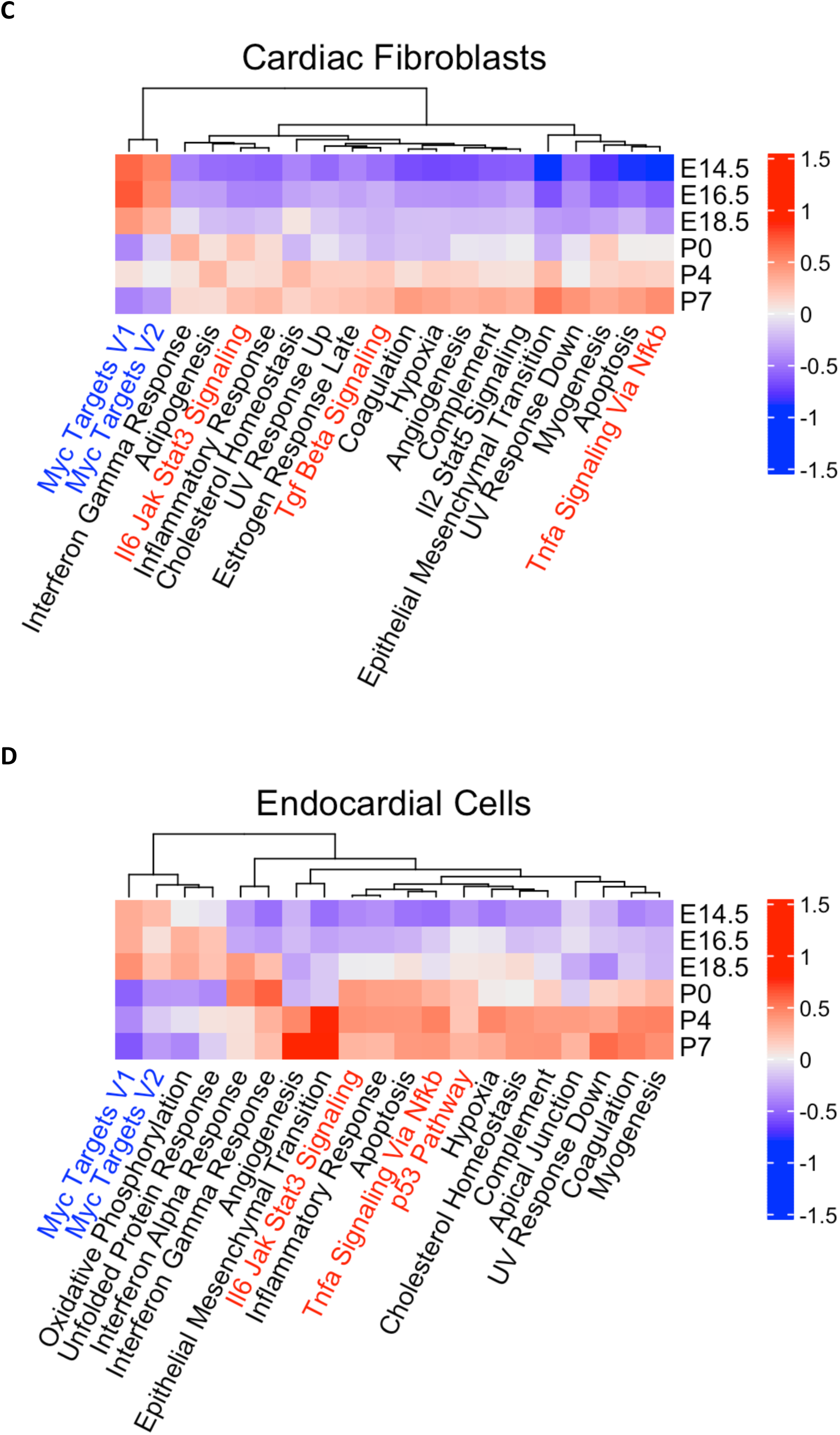
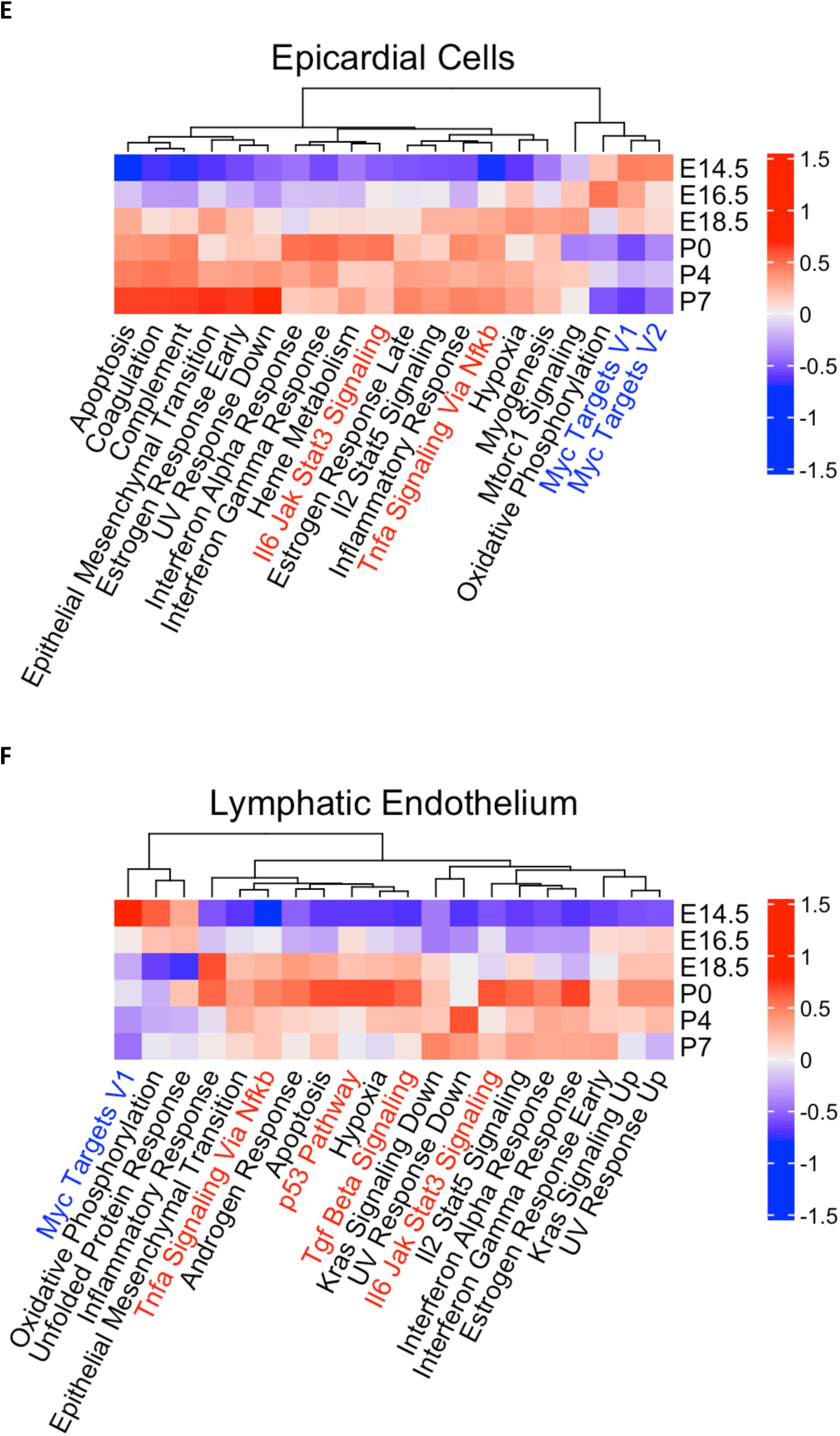
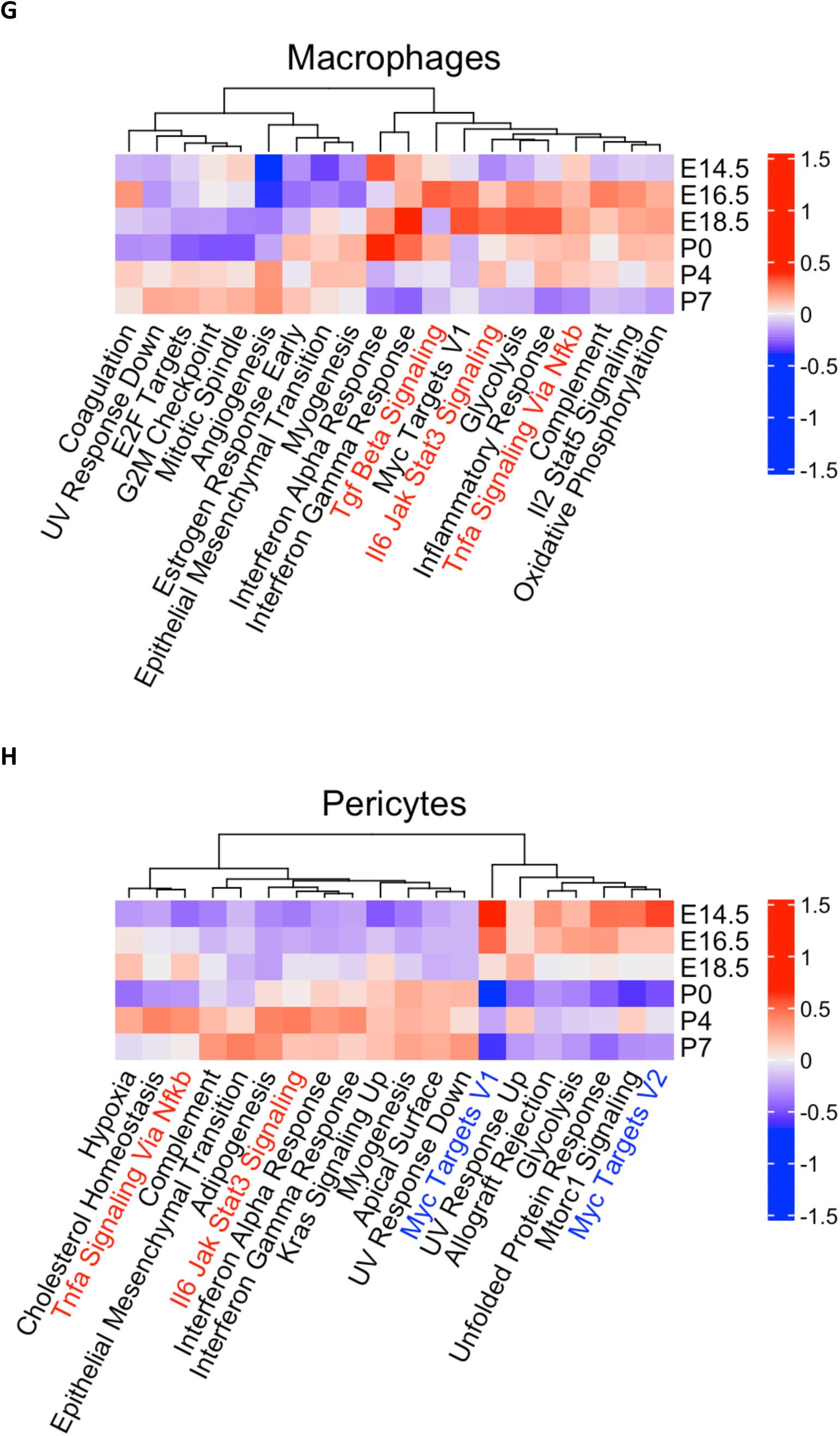
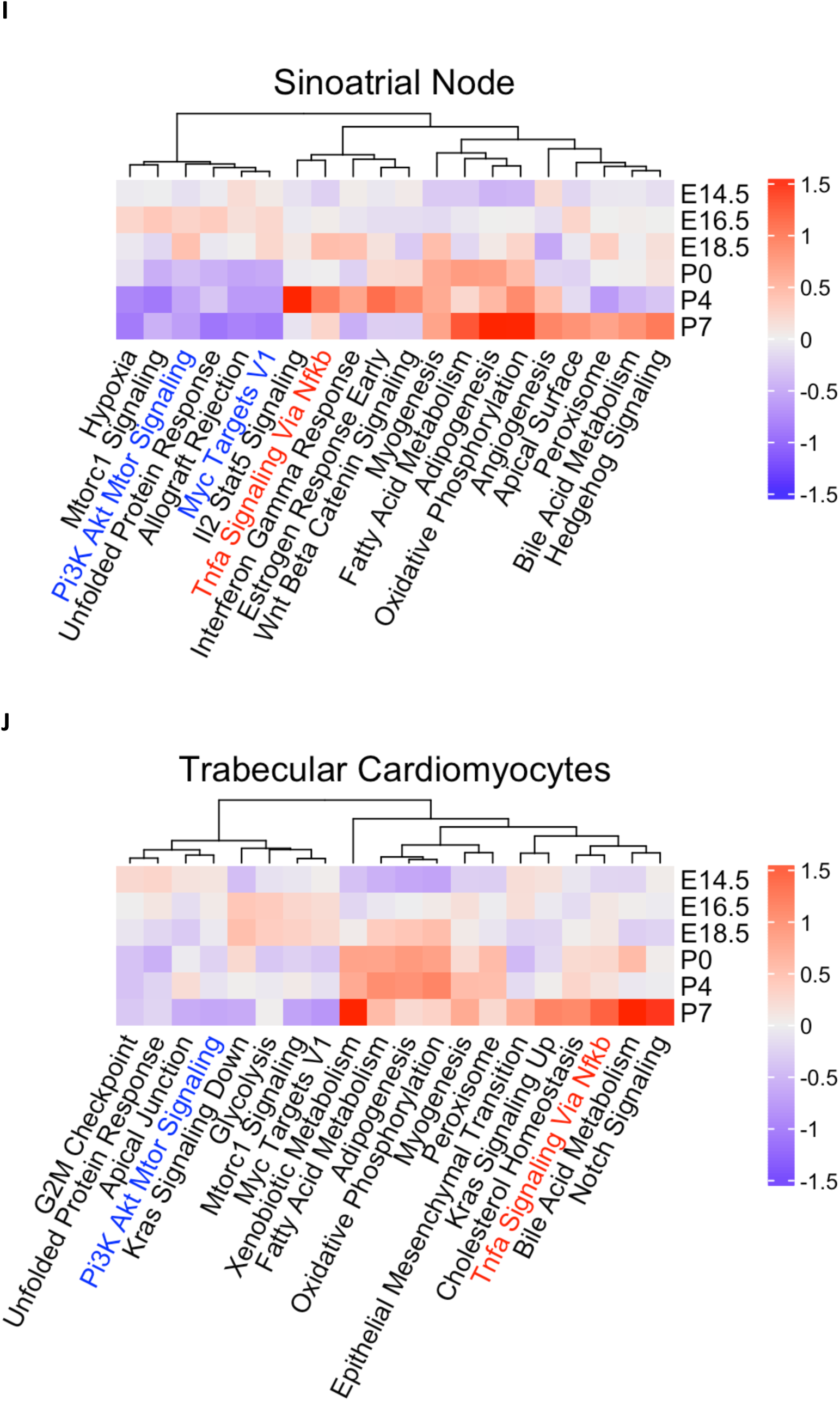
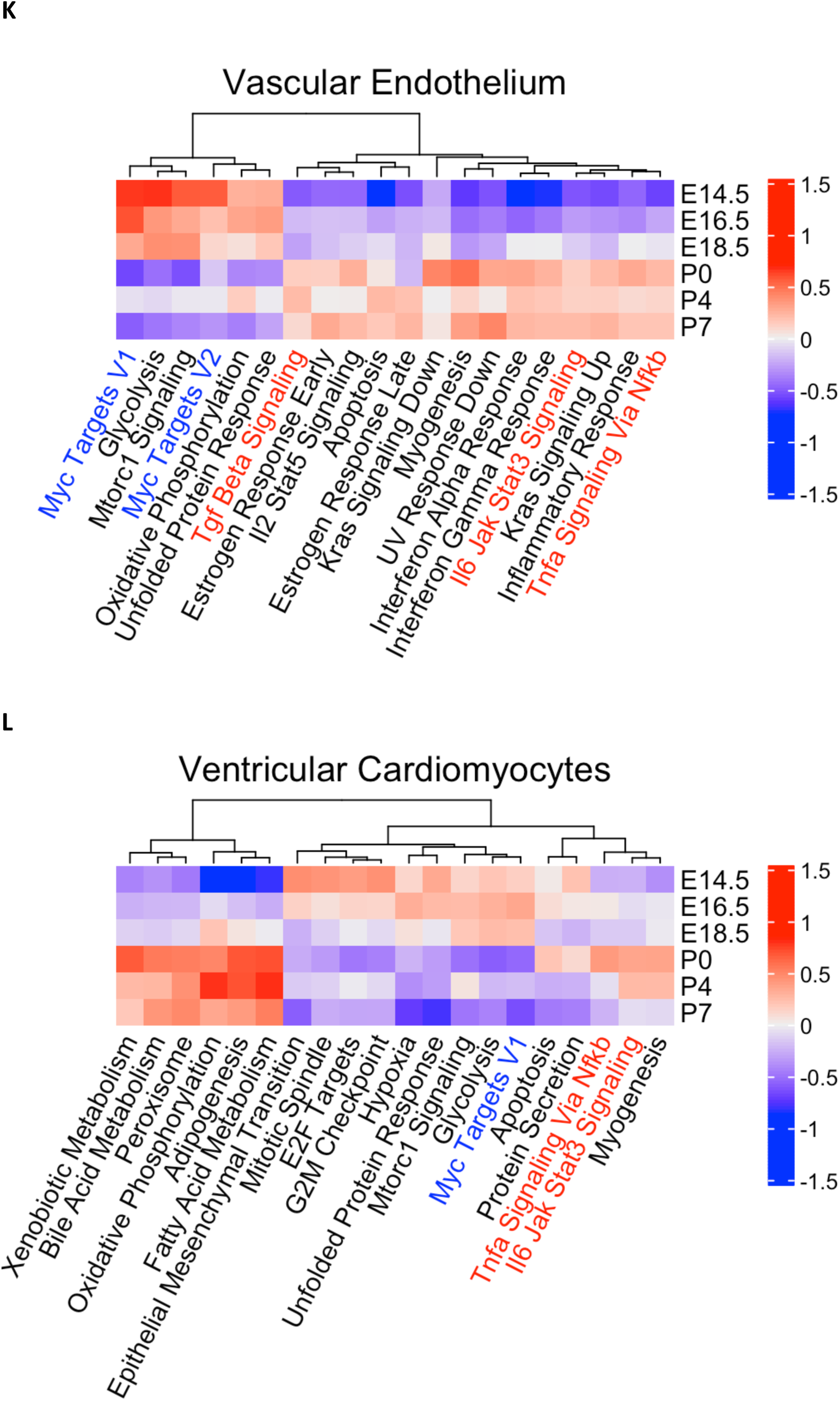
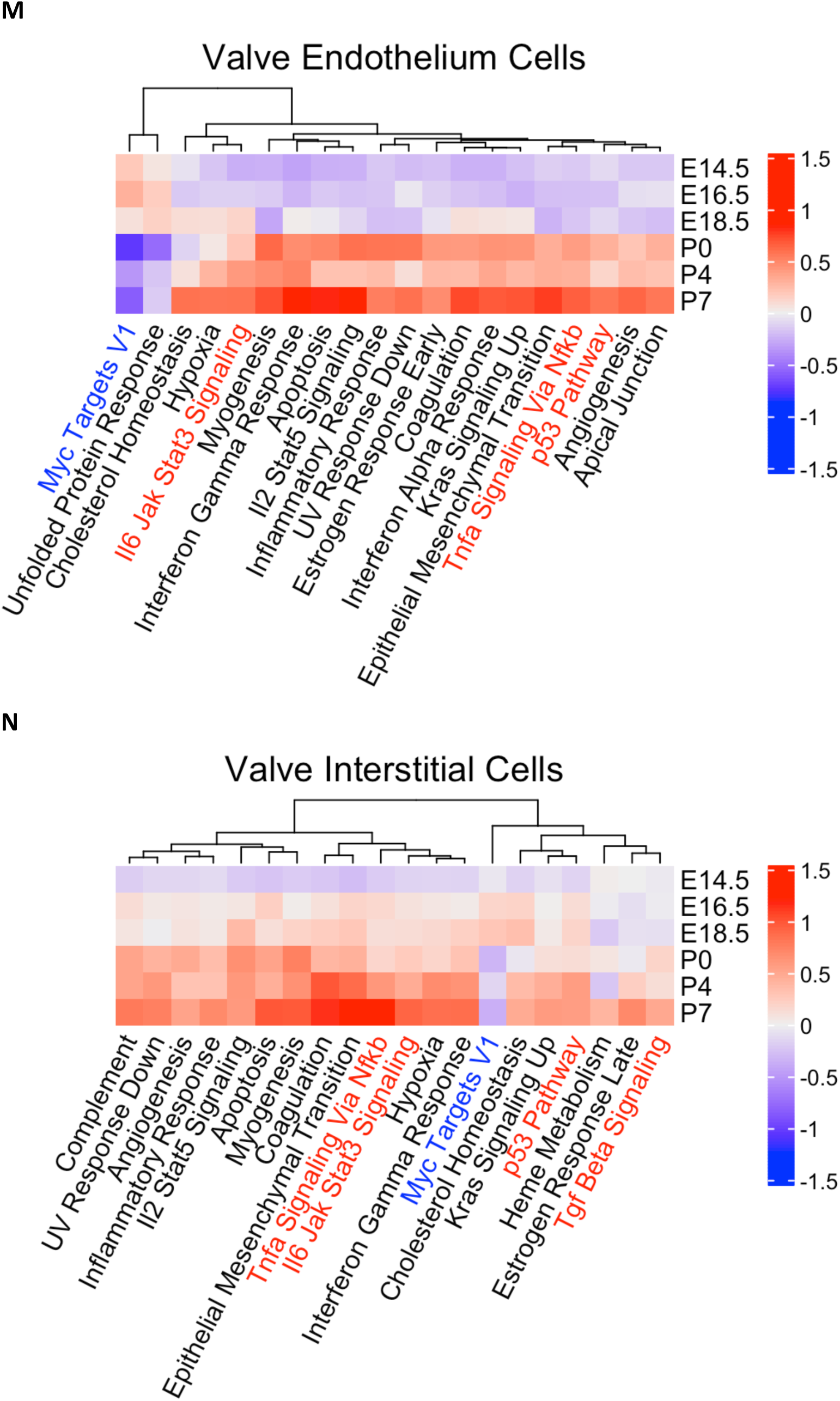
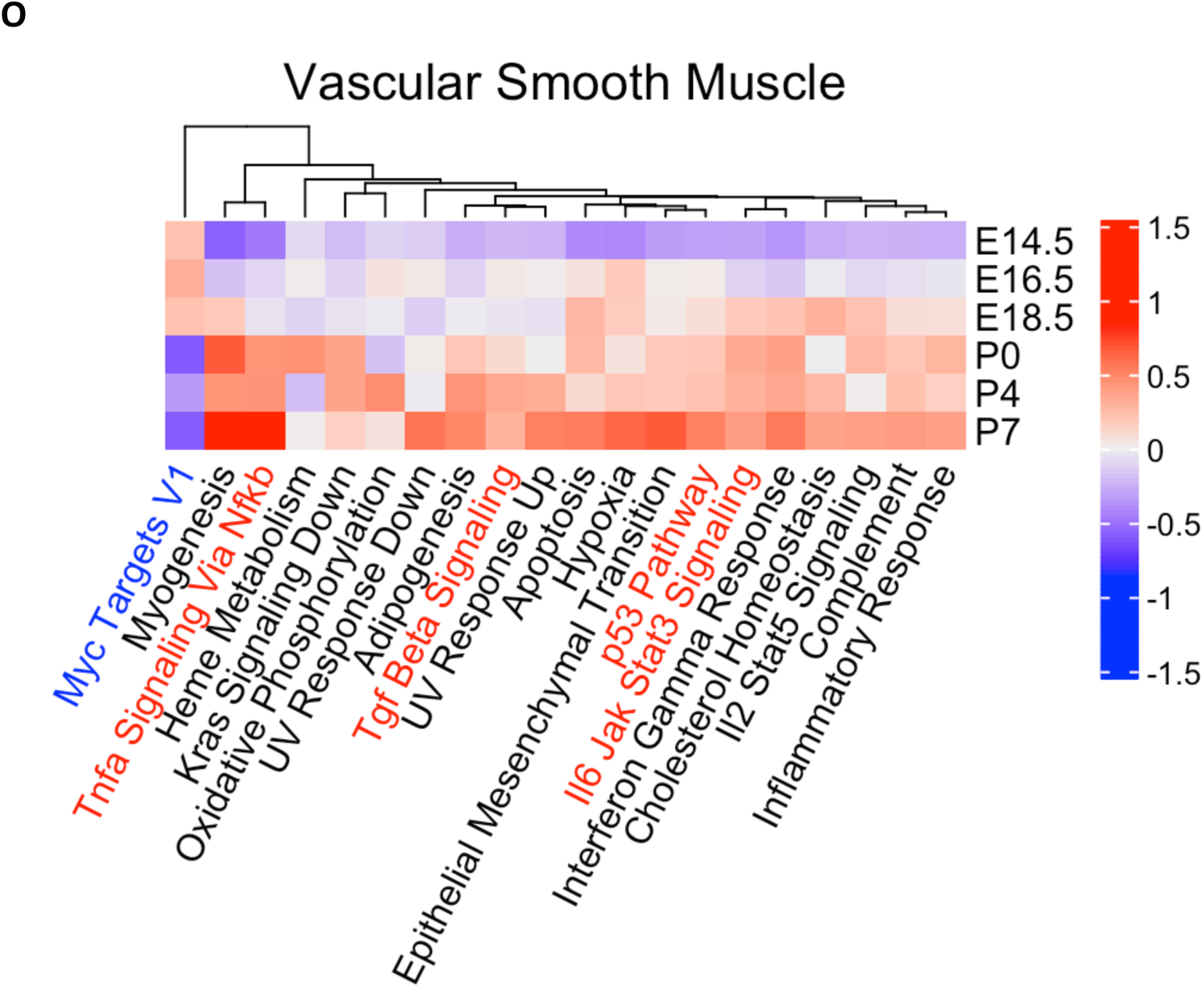
decoupleR Hallmark pathway activity heatmaps as in Figure 5 for all analysed cell types.

**Figure S3:**
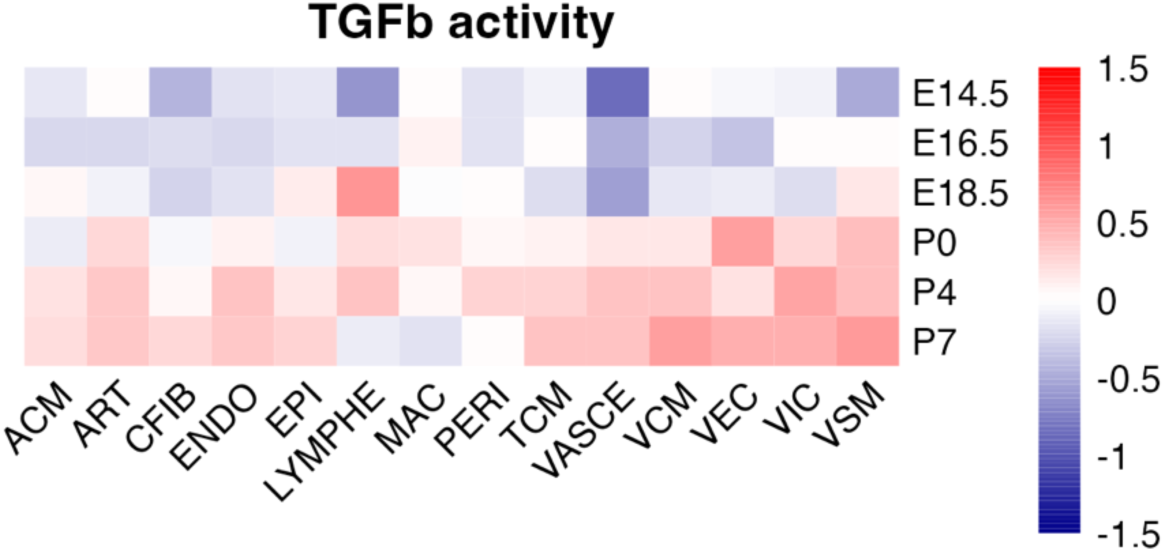
Compiled heatmap of TGFβ pathway activity generated by decoupleR using the PROGENy database. TGFβ pathway increases with developmental time across most cardiac cell types.

**Figure S4:**
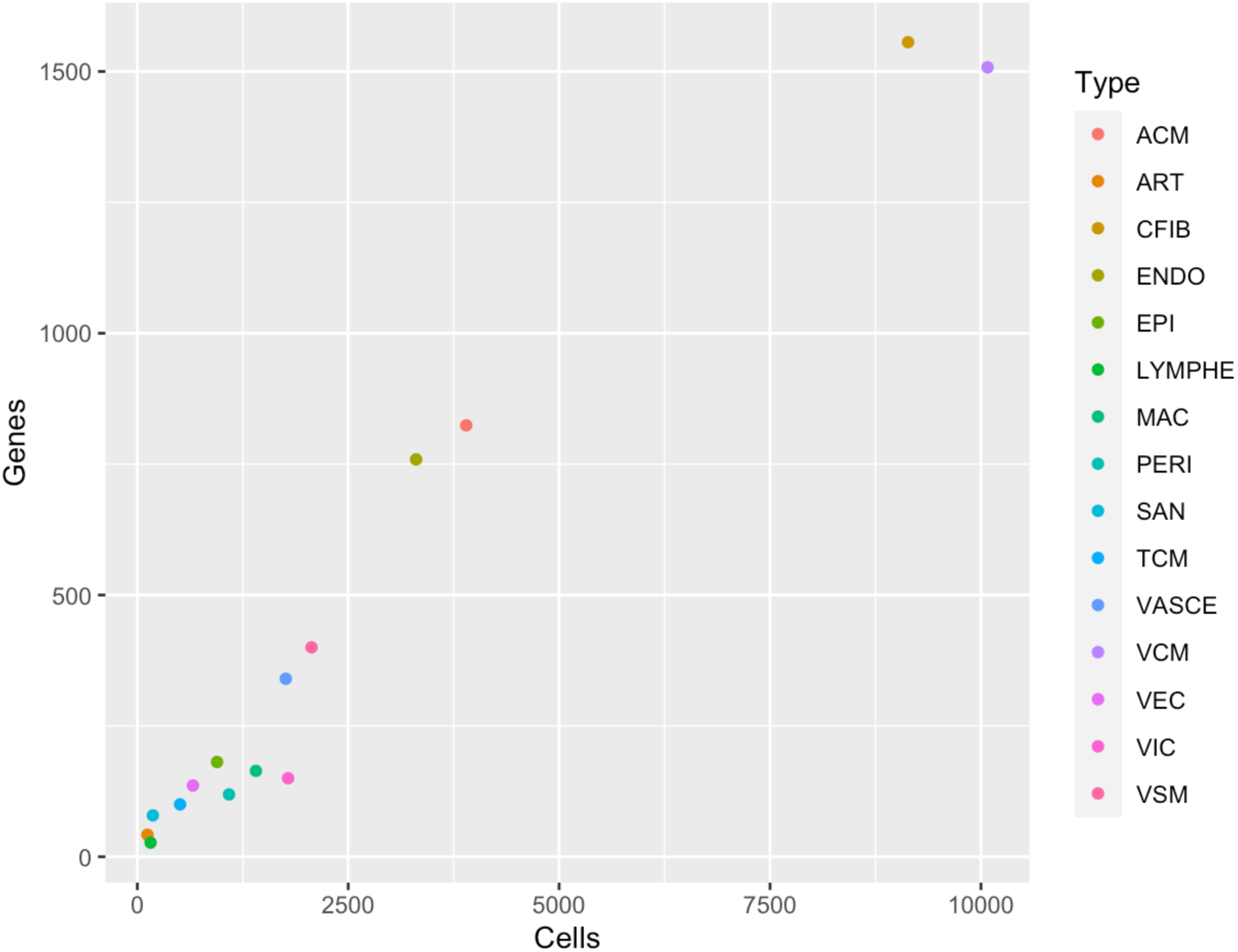
Scatter plot showing positive correlation between the number of cells in a cell type and the number of time-varying genes identified by psupertime.

**Figure S5:**
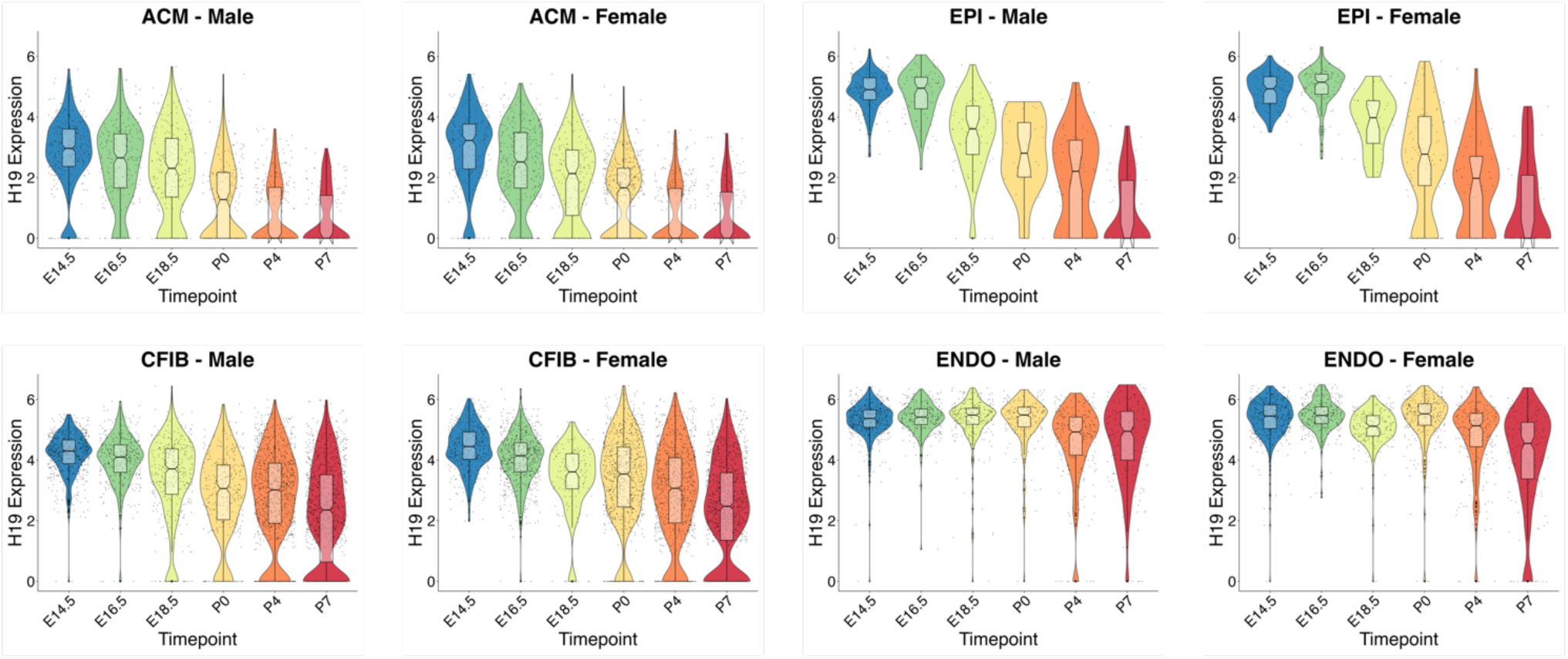
Seurat violin plots of H19 gene expression as in Figure 7A for male and female samples only. The expression pattern of H19 does not appear to be sex-specific.

**Figure S6:**
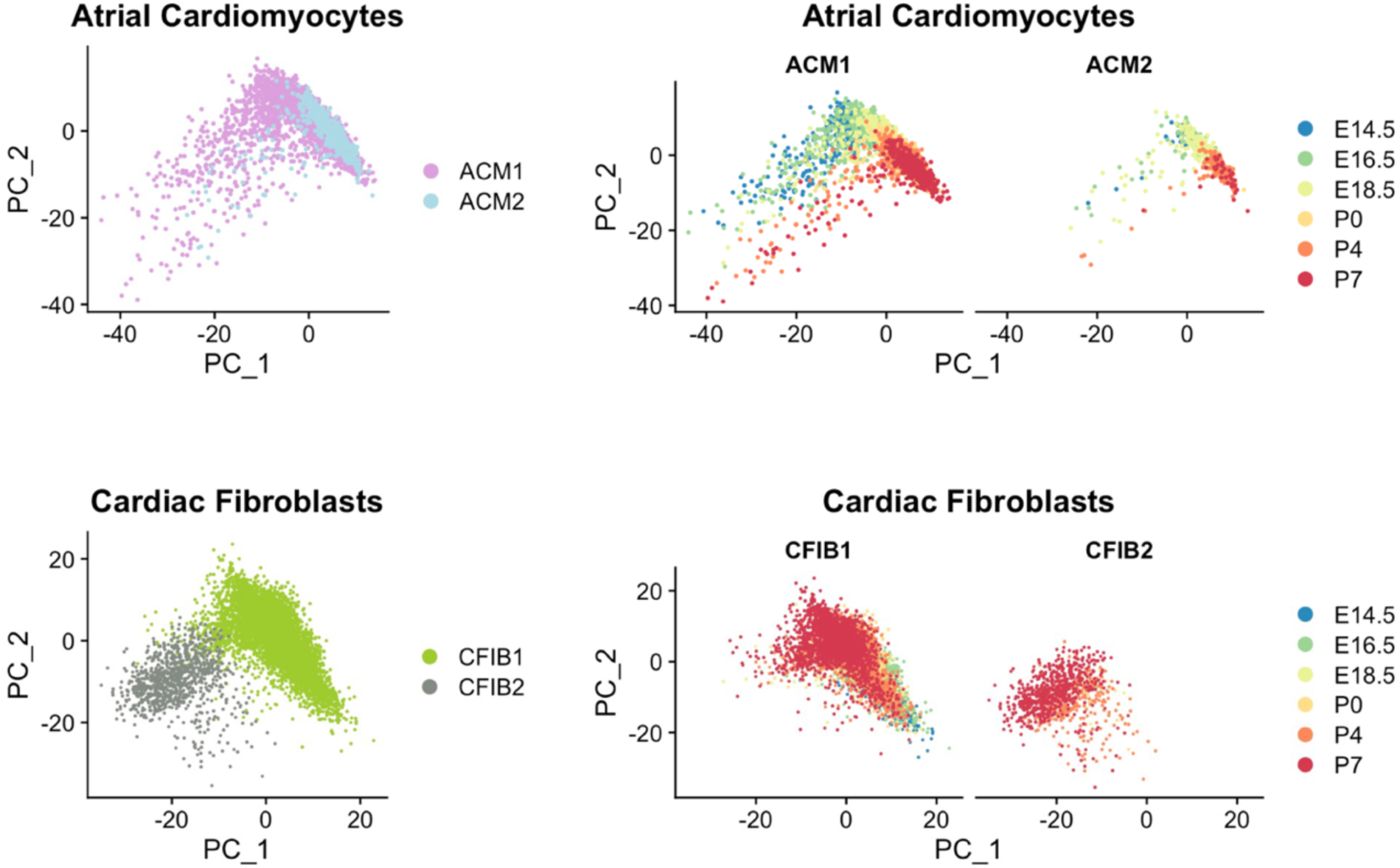
PCA plots of all atrial cardiomyocytes and all cardiac fibroblasts. The ACM1 and ACM2 clusters combine to form a continuous trajectory, while CFIB1 and CFIB2 appear to be divergent.

## Supplementary Tables

**Table S1: Metadata information for all cells in final single-cell dataset.**

Fastq files for E14.5 samples were previously published (Wünnemann et al.) and reanalyzed from raw data using the same pipeline used for all other samples. Six replicates were used per time point for a total of 36 mice as indicated in the Sample column (ex. E14.5_r1). nCount: number of unique molecules (UMIs), nFeature: number of genes per cell, SCT_snn_res: cell cluster number for various resolutions.

**Table S2: Cell-type marker genes for all cell types in the global cell clustering.**

Marker genes that identify cell types were identified using the Seurat function FindMarkers(). avg_log2FC: log2 fold change, pval: p-value, p_val_adjusted: Benjamini-Hochberg adjusted p-value, pct.1: percent of observations in given cell types, pct.2: percent of observations in all other cell types.

**Table S3:** Foetal and postnatal regulon specificity scores (RSS) from SCENIC analysis ranging from 0 (least specific) to 1 (most specific) for all regulons and all cell types.

**Table S4: Differentially expressed genes across pseudotime for all cell types.** psupertime results for all individual cell types analysed. abs_beta: absolute beta coefficient, a score >0 indicates significant variance across time. *H19* was among or the highest scoring in numerous cell types.

**Table S5: Results from enrichment analysis using enrichR against the MGI Mammalian phenotype level 4 2021 database.**

Genes with significant differential expression across pseudotime per psupertime (abs_beta>0 from Table S4) in more than 2 cell types were tested as one gene list. The p-values were computed using Fisher’s exact test, while the combined enrichment scores were calculated by multiplying the log of the p-value with the odds ratio.

## References

Aibar, S., González-Blas, C.B., Moerman, T., Huynh-Thu, V.A., Imrichova, H., Hulselmans, G., Rambow, F., Marine, J.-C., Geurts, P., Aerts, J., van den Oord, J., Atak, Z.K., Wouters, J., Aerts, S., 2017. SCENIC: single-cell regulatory network inference and clustering. Nat. Methods 14, 1083–1086. 10.1038/nmeth.446310.1161/CIRCRESAHA.116.308139

Anderson, R.H., Spicer, D.E., Brown, N.A., Mohun, T.J., 2014. The development of septation in the four-chambered heart. Anat. Rec. Hoboken NJ 2007 297, 1414–1429. 10.1002/ar.22949

Asp, M., Giacomello, S., Larsson, L., Wu, C., Fürth, D., Qian, X., Wärdell, E., Custodio, J., Reimegård, J., Salmén, F., Österholm, C., Ståhl, P.L., Sundström, E., Åkesson, E., Bergmann, O., Bienko, M., Månsson-Broberg, A., Nilsson, M., Sylvén, C., Lundeberg, J., 2019. A Spatiotemporal Organ-Wide Gene Expression and Cell Atlas of the Developing Human Heart. Cell 179, 1647–1660.e19. 10.1016/j.cell.2019.11.025

Badia-I-Mompel, P., Vélez Santiago, J., Braunger, J., Geiss, C., Dimitrov, D., Müller-Dott, S., Taus, P., Dugourd, A., Holland, C.H., Ramirez Flores, R.O., Saez-Rodriguez, J., 2022. decoupleR: ensemble of computational methods to infer biological activities from omics data. Bioinforma. Adv. 2, vbac016. 10.1093/bioadv/vbac016

Banerjee, I., Fuseler, J.W., Price, R.L., Borg, T.K., Baudino, T.A., 2007. Determination of cell types and numbers during cardiac development in the neonatal and adult rat and mouse. Am. J. Physiol. Heart Circ. Physiol. 293, H1883–1891. 10.1152/ajpheart.00514.2007

Bedada, F.B., Chan, S.S.-K., Metzger, S.K., Zhang, L., Zhang, J., Garry, D.J., Kamp, T.J., Kyba, M., Metzger, J.M., 2014. Acquisition of a quantitative, stoichiometrically conserved ratiometric marker of maturation status in stem cell-derived cardiac myocytes. Stem Cell Rep. 3, 594–605. 10.1016/j.stemcr.2014.07.012

Bliek, J., Terhal, P., van den Bogaard, M.-J., Maas, S., Hamel, B., Salieb-Beugelaar, G., Simon, M., Letteboer, T., van der Smagt, J., Kroes, H., Mannens, M., 2006. Hypomethylation of the H19 gene causes not only Silver-Russell syndrome (SRS) but also isolated asymmetry or an SRS-like phenotype. Am. J. Hum. Genet. 78, 604–614. 10.1086/502981

Bruneau, B.G., 2013. Signaling and transcriptional networks in heart development and regeneration. Cold Spring Harb. Perspect. Biol. 5, a008292. 10.1101/cshperspect.a008292

Brunkow, M.E., Tilghman, S.M., 1991. Ectopic expression of the H19 gene in mice causes prenatal lethality. Genes Dev. 5, 1092–1101. 10.1101/gad.5.6.1092

Buckingham, M., Meilhac, S., Zaffran, S., 2005. Building the mammalian heart from two sources of myocardial cells. Nat. Rev. Genet. 6, 826–835. 10.1038/nrg1710

Cai, C.-L., Liang, X., Shi, Y., Chu, P.-H., Pfaff, S.L., Chen, J., Evans, S., 2003. Isl1 identifies a cardiac progenitor population that proliferates prior to differentiation and contributes a majority of cells to the heart. Dev. Cell 5, 877–889. 10.1016/s1534-5807(03)00363-0

Castanza, A.S., Recla, J.M., Eby, D., Thorvaldsdóttir, H., Bult, C.J., Mesirov, J.P., 2023. Extending support for mouse data in the Molecular Signatures Database (MSigDB). Nat. Methods 20, 1619–1620. 10.1038/s41592-023-02014-7

Chen, E.Y., Tan, C.M., Kou, Y., Duan, Q., Wang, Z., Meirelles, G.V., Clark, N.R., Ma’ayan, A., 2013. Enrichr: interactive and collaborative HTML5 gene list enrichment analysis tool. BMC Bioinformatics 14, 128. 10.1186/1471-2105-14-128

Chiplunkar, A.R., Lung, T.K., Alhashem, Y., Koppenhaver, B.A., Salloum, F.N., Kukreja, R.C., Haar, J.L., Lloyd, J.A., 2013. Krüppel-like factor 2 is required for normal mouse cardiac development. PloS One 8, e54891. 10.1371/journal.pone.0054891

Cohen, A.W., Park, D.S., Woodman, S.E., Williams, T.M., Chandra, M., Shirani, J., Pereira de Souza, A., Kitsis, R.N., Russell, R.G., Weiss, L.M., Tang, B., Jelicks, L.A., Factor, S.M., Shtutin, V., Tanowitz, H.B., Lisanti, M.P., 2003. Caveolin-1 null mice develop cardiac hypertrophy with hyperactivation of p42/44 MAP kinase in cardiac fibroblasts. Am. J. Physiol. Cell Physiol. 284, C457–474. 10.1152/ajpcell.00380.2002

Cook, D.P., Vanderhyden, B.C., 2020. Context specificity of the EMT transcriptional response. Nat. Commun. 11, 2142. 10.1038/s41467-020-16066-2

Cui, Y., Zheng, Y., Liu, X., Yan, L., Fan, X., Yong, J., Hu, Y., Dong, J., Li, Q., Wu, X., Gao, S., Li, J., Wen, L., Qiao, J., Tang, F., 2019. Single-Cell Transcriptome Analysis Maps the Developmental Track of the Human Heart. Cell Rep. 26, 1934–1950.e5. 10.1016/j.celrep.2019.01.079

Dawson, J.A., Kamlin, C.O.F., Vento, M., Wong, C., Cole, T.J., Donath, S.M., Davis, P.G., Morley, C.J., 2010. Defining the reference range for oxygen saturation for infants after birth. Pediatrics 125, e1340–1347. 10.1542/peds.2009-1510

de Soysa, T.Y., Ranade, S.S., Okawa, S., Ravichandran, S., Huang, Y., Salunga, H.T., Schricker, A., Del Sol, A., Gifford, C.A., Srivastava, D., 2019. Single-cell analysis of cardiogenesis reveals basis for organ-level developmental defects. Nature 572, 120–124. 10.1038/s41586-019-1414-x

del Monte, G., Casanova, J.C., Guadix, J.A., MacGrogan, D., Burch, J.B.E., Pérez-Pomares, J.M., de la Pompa, J.L., 2011. Differential Notch signaling in the epicardium is required for cardiac inflow development and coronary vessel morphogenesis. Circ. Res. 108, 824–836. 10.1161/CIRCRESAHA.110.229062

DeLaughter, D.M., Bick, A.G., Wakimoto, H., McKean, D., Gorham, J.M., Kathiriya, I.S., Hinson, J.T., Homsy, J., Gray, J., Pu, W., Bruneau, B.G., Seidman, J.G., Seidman, C.E., 2016. Single-Cell Resolution of Temporal Gene Expression during Heart Development. Dev. Cell 39, 480–490. 10.1016/j.devcel.2016.10.001

Dhingra, S., Sharma, A.K., Arora, R.C., Slezak, J., Singal, P.K., 2009. IL-10 attenuates TNF-alpha-induced NF kappaB pathway activation and cardiomyocyte apoptosis. Cardiovasc. Res. 82, 59–66. 10.1093/cvr/cvp040

Dobaczewski, M., Chen, W., Frangogiannis, N.G., 2011. Transforming growth factor (TGF)-β signaling in cardiac remodeling. J. Mol. Cell. Cardiol. 51, 600–606. 10.1016/j.yjmcc.2010.10.033

Durinck, S., Moreau, Y., Kasprzyk, A., Davis, S., De Moor, B., Brazma, A., Huber, W., 2005. BioMart and Bioconductor: a powerful link between biological databases and microarray data analysis. Bioinforma. Oxf. Engl. 21, 3439–3440. 10.1093/bioinformatics/bti525

Duronio, R.J., Xiong, Y., 2013. Signaling pathways that control cell proliferation. Cold Spring Harb. Perspect. Biol. 5, a008904. 10.1101/cshperspect.a008904

Duyao, M.P., Buckler, A.J., Sonenshein, G.E., 1990. Interaction of an NF-kappa B-like factor with a site upstream of the c-myc promoter. Proc. Natl. Acad. Sci. U. S. A. 87, 4727–4731. 10.1073/pnas.87.12.4727

Eferl, R., Sibilia, M., Hilberg, F., Fuchsbichler, A., Kufferath, I., Guertl, B., Zenz, R., Wagner, E.F., Zatloukal, K., 1999. Functions of c-Jun in liver and heart development. J. Cell Biol. 145, 1049–1061. 10.1083/jcb.145.5.1049

Feng, W., Bais, A., He, H., Rios, C., Jiang, S., Xu, J., Chang, C., Kostka, D., Li, G., 2022. Single-cell transcriptomic analysis identifies murine heart molecular features at embryonic and neonatal stages. Nat. Commun. 13, 7960. 10.1038/s41467-022-35691-7

Feng, W., Chen, L., Nguyen, P.K., Wu, S.M., Li, G., 2019. Single Cell Analysis of Endothelial Cells Identified Organ-Specific Molecular Signatures and Heart-Specific Cell Populations and Molecular Features. Front. Cardiovasc. Med. 6, 165. 10.3389/fcvm.2019.00165

Ferdous, A., Caprioli, A., Iacovino, M., Martin, C.M., Morris, J., Richardson, J.A., Latif, S., Hammer, R.E., Harvey, R.P., Olson, E.N., Kyba, M., Garry, D.J., 2009. Nkx2-5 transactivates the Ets-related protein 71 gene and specifies an endothelial/endocardial fate in the developing embryo. Proc. Natl. Acad. Sci. U. S. A. 106, 814–819. 10.1073/pnas.0807583106

Feulner, L., van Vliet, P.P., Puceat, M., Andelfinger, G., 2022. Endocardial Regulation of Cardiac Development. J. Cardiovasc. Dev. Dis. 9, 122. 10.3390/jcdd9050122

Filipczyk, A.A., Passier, R., Rochat, A., Mummery, C.L., 2007. Regulation of cardiomyocyte differentiation of embryonic stem cells by extracellular signalling. Cell. Mol. Life Sci. CMLS 64, 704–718. 10.1007/s00018-007-6523-2

Forte, E., Skelly, D.A., Chen, M., Daigle, S., Morelli, K.A., Hon, O., Philip, V.M., Costa, M.W., Rosenthal, N.A., Furtado, M.B., 2020. Dynamic Interstitial Cell Response during Myocardial Infarction Predicts Resilience to Rupture in Genetically Diverse Mice. Cell Rep. 30, 3149–3163.e6. 10.1016/j.celrep.2020.02.008

Frangogiannis, N.G., 2022. Transforming growth factor-β in myocardial disease. Nat. Rev. Cardiol. 19, 435–455. 10.1038/s41569-021-00646-w

Gabory, A., Jammes, H., Dandolo, L., 2010. The H19 locus: role of an imprinted non-coding RNA in growth and development. BioEssays News Rev. Mol. Cell. Dev. Biol. 32, 473–480. 10.1002/bies.200900170

Ghanim, M., Rossignol, S., Delobel, B., Irving, M., Miller, O., Devisme, L., Plennevaux, J.-L., Lucidarme-Rossi, S., Manouvrier, S., Salah, A., Chivu, O., Netchine, I., Vincent-Delorme, C., 2013. Possible association between complex congenital heart defects and 11p15 hypomethylation in three patients with severe Silver-Russell syndrome. Am. J. Med. Genet. A. 161A, 572–577. 10.1002/ajmg.a.35691

Gierahn, T.M., Wadsworth, M.H., Hughes, T.K., Bryson, B.D., Butler, A., Satija, R., Fortune, S., Love, J.C., Shalek, A.K., 2017. Seq-Well: portable, low-cost RNA sequencing of single cells at high throughput. Nat. Methods 14, 395–398. 10.1038/nmeth.4179

Goddard, L.M., Duchemin, A.-L., Ramalingan, H., Wu, B., Chen, M., Bamezai, S., Yang, J., Li, L., Morley, M.P., Wang, T., Scherrer-Crosbie, M., Frank, D.B., Engleka, K.A., Jameson, S.C., Morrisey, E.E., Carroll, T.J., Zhou, B., Vermot, J., Kahn, M.L., 2017. Hemodynamic Forces Sculpt Developing Heart Valves through a KLF2-WNT9B Paracrine Signaling Axis. Dev. Cell 43, 274–289.e5. 10.1016/j.devcel.2017.09.023

Goodyer, W.R., Beyersdorf, B.M., Paik, D.T., Tian, L., Li, G., Buikema, J.W., Chirikian, O., Choi, S., Venkatraman, S., Adams, E.L., Tessier-Lavigne, M., Wu, J.C., Wu, S.M., 2019. Transcriptomic Profiling of the Developing Cardiac Conduction System at Single-Cell Resolution. Circ. Res. 125, 379–397. 10.1161/CIRCRESAHA.118.314578

Greenwood, R.D., Somer, A., Rosenthal, A., Craenen, J., Nadas, A.S., 1977. Cardiovascular abnormalities in the Beckwith-Wiedemann syndrome. Am. J. Dis. Child. 1960 131, 293–294. 10.1001/archpedi.1977.02120160047007

Hadji, F., Boulanger, M.-C., Guay, S.-P., Gaudreault, N., Amellah, S., Mkannez, G., Bouchareb, R., Marchand, J.T., Nsaibia, M.J., Guauque-Olarte, S., Pibarot, P., Bouchard, L., Bossé, Y., Mathieu, P., 2016. Altered DNA Methylation of Long Noncoding RNA H19 in Calcific Aortic Valve Disease Promotes Mineralization by Silencing NOTCH1. Circulation 134, 1848–1862. 10.1161/CIRCULATIONAHA.116.023116

Hafemeister, C., Satija, R., 2019. Normalization and variance stabilization of single-cell RNA-seq data using regularized negative binomial regression. Genome Biol. 20, 296. 10.1186/s13059-019-1874-1

Hirano, T., Ishihara, K., Hibi, M., 2000. Roles of STAT3 in mediating the cell growth, differentiation and survival signals relayed through the IL-6 family of cytokine receptors. Oncogene 19, 2548–2556. 10.1038/sj.onc.1203551

Hortells, L., Valiente-Alandi, I., Thomas, Z.M., Agnew, E.J., Schnell, D.J., York, A.J., Vagnozzi, R.J., Meyer, E.C., Molkentin, J.D., Yutzey, K.E., 2020. A specialized population of Periostin-expressing cardiac fibroblasts contributes to postnatal cardiomyocyte maturation and innervation. Proc. Natl. Acad. Sci. U. S. A. 117, 21469–21479. 10.1073/pnas.2009119117

Houweling, A.C., van Borren, M.M., Moorman, A.F.M., Christoffels, V.M., 2005. Expression and regulation of the atrial natriuretic factor encoding gene Nppa during development and disease. Cardiovasc. Res. 67, 583–593. 10.1016/j.cardiores.2005.06.013

Hu, P., Liu, J., Zhao, J., Wilkins, B.J., Lupino, K., Wu, H., Pei, L., 2018. Single-nucleus transcriptomic survey of cell diversity and functional maturation in postnatal mammalian hearts. Genes Dev. 32, 1344–1357. 10.1101/gad.316802.118

Ilya Korsunsky, Aparna Nathan, Nghia Millard, Soumya Raychaudhuri, 2019. Presto scales Wilcoxon and auROC analyses to millions of observations. bioRxiv 653253. 10.1101/653253

Kannan, S., Farid, M., Lin, B.L., Miyamoto, M., Kwon, C., 2021. Transcriptomic entropy benchmarks stem cell-derived cardiomyocyte maturation against endogenous tissue at single cell level. PLoS Comput. Biol. 17, e1009305. 10.1371/journal.pcbi.1009305

Kiuchi, N., Nakajima, K., Ichiba, M., Fukada, T., Narimatsu, M., Mizuno, K., Hibi, M., Hirano, T., 1999. STAT3 is required for the gp130-mediated full activation of the c-myc gene. J. Exp. Med. 189, 63–73. 10.1084/jem.189.1.63

Kodama, H., Fukuda, K., Pan, J., Makino, S., Baba, A., Hori, S., Ogawa, S., 1997. Leukemia inhibitory factor, a potent cardiac hypertrophic cytokine, activates the JAK/STAT pathway in rat cardiomyocytes. Circ. Res. 81, 656–663. 10.1161/01.res.81.5.656

Koenig, A.L., Shchukina, I., Amrute, J., Andhey, P.S., Zaitsev, K., Lai, L., Bajpai, G., Bredemeyer, A., Smith, G., Jones, C., Terrebonne, E., Rentschler, S.L., Artyomov, M.N., Lavine, K.J., 2022. Single-cell transcriptomics reveals cell-type-specific diversification in human heart failure. Nat. Cardiovasc. Res. 1, 263–280. 10.1038/s44161-022-00028-6

Kuleshov, M.V., Jones, M.R., Rouillard, A.D., Fernandez, N.F., Duan, Q., Wang, Z., Koplev, S., Jenkins, S.L., Jagodnik, K.M., Lachmann, A., McDermott, M.G., Monteiro, C.D., Gundersen, G.W., Ma’ayan, A., 2016. Enrichr: a comprehensive gene set enrichment analysis web server 2016 update. Nucleic Acids Res. 44, W90–97. 10.1093/nar/gkw377

Kuwahara, K., Nishikimi, T., Nakao, K., 2012. Transcriptional regulation of the fetal cardiac gene program. J. Pharmacol. Sci. 119, 198–203. 10.1254/jphs.12r04cp

L Bingle, H Armes, DJ Williams, O Gianfrancesco, Md M K Chowdhury, R Drapkin, C D Bingle, 2020. Expression and function of murine WFDC2 in the respiratory tract. bioRxiv 2020.05.05.079293. 10.1101/2020.05.05.079293

Lai, L., Leone, T.C., Zechner, C., Schaeffer, P.J., Kelly, S.M., Flanagan, D.P., Medeiros, D.M., Kovacs, A., Kelly, D.P., 2008. Transcriptional coactivators PGC-1alpha and PGC-lbeta control overlapping programs required for perinatal maturation of the heart. Genes Dev. 22, 1948–1961. 10.1101/gad.1661708

Li, G., Xu, A., Sim, S., Priest, J.R., Tian, X., Khan, T., Quertermous, T., Zhou, B., Tsao, P.S., Quake, S.R., Wu, S.M., 2016. Transcriptomic Profiling Maps Anatomically Patterned Subpopulations among Single Embryonic Cardiac Cells. Dev. Cell 39, 491–507. 10.1016/j.devcel.2016.10.014

Li, Z., Solomonidis, E.G., Meloni, M., Taylor, R.S., Duffin, R., Dobie, R., Magalhaes, M.S., Henderson, B.E.P., Louwe, P.A., D’Amico, G., Hodivala-Dilke, K.M., Shah, A.M., Mills, N.L., Simons, B.D., Gray, G.A., Henderson, N.C., Baker, A.H., Brittan, M., 2019. Single-cell transcriptome analyses reveal novel targets modulating cardiac neovascularization by resident endothelial cells following myocardial infarction. Eur. Heart J. 40, 2507–2520. 10.1093/eurheartj/ehz305

Lints, T.J., Parsons, L.M., Hartley, L., Lyons, I., Harvey, R.P., 1993. Nkx-2.5: a novel murine homeobox gene expressed in early heart progenitor cells and their myogenic descendants. Dev. Camb. Engl. 119, 419–431.

Litviňuková, M., Talavera-López, C., Maatz, H., Reichart, D., Worth, C.L., Lindberg, E.L., Kanda, M., Polanski, K., Heinig, M., Lee, M., Nadelmann, E.R., Roberts, K., Tuck, L., Fasouli, E.S., DeLaughter, D.M., McDonough, B., Wakimoto, H., Gorham, J.M., Samari, S., Mahbubani, K.T., Saeb-Parsy, K., Patone, G., Boyle, J.J., Zhang, Hongbo, Zhang, Hao, Viveiros, A., Oudit, G.Y., Bayraktar, O.A., Seidman, J.G., Seidman, C.E., Noseda, M., Hubner, N., Teichmann, S.A., 2020. Cells of the adult human heart. Nature 588, 466–472. 10.1038/s41586-020-2797-4

Liu, L., Brown, D., McKee, M., Lebrasseur, N.K., Yang, D., Albrecht, K.H., Ravid, K., Pilch, P.F., 2008. Deletion of Cavin/PTRF causes global loss of caveolae, dyslipidemia, and glucose intolerance. Cell Metab. 8, 310–317. 10.1016/j.cmet.2008.07.008

Lui, J.C., Finkielstain, G.P., Barnes, K.M., Baron, J., 2008. An imprinted gene network that controls mammalian somatic growth is down-regulated during postnatal growth deceleration in multiple organs. Am. J. Physiol. Regul. Integr. Comp. Physiol. 295, R189–196. 10.1152/ajpregu.00182.2008

Lun, A.T.L., Riesenfeld, S., Andrews, T., Dao, T.P., Gomes, T., participants in the 1st Human Cell Atlas Jamboree, Marioni, J.C., 2019. EmptyDrops: distinguishing cells from empty droplets in droplet-based single-cell RNA sequencing data. Genome Biol. 20, 63. 10.1186/s13059-019-1662-y

Luo, K., Stroschein, S.L., Wang, W., Chen, D., Martens, E., Zhou, S., Zhou, Q., 1999. The Ski oncoprotein interacts with the Smad proteins to repress TGFbeta signaling. Genes Dev. 13, 2196–2206. 10.1101/gad.13.17.2196

Macnair, W., Gupta, R., Claassen, M., 2022. psupertime: supervised pseudotime analysis for time-series single-cell RNA-seq data. Bioinforma. Oxf. Engl. 38, i290–i298. 10.1093/bioinformatics/btac227

Macosko, E.Z., Basu, A., Satija, R., Nemesh, J., Shekhar, K., Goldman, M., Tirosh, I., Bialas, A.R., Kamitaki, N., Martersteck, E.M., Trombetta, J.J., Weitz, D.A., Sanes, J.R., Shalek, A.K., Regev, A., McCarroll, S.A., 2015. Highly Parallel Genome-wide Expression Profiling of Individual Cells Using Nanoliter Droplets. Cell 161, 1202–1214. 10.1016/j.cell.2015.05.002

Mak, T.W., Hauck, L., Grothe, D., Billia, F., 2017. p53 regulates the cardiac transcriptome. Proc. Natl. Acad. Sci. U. S. A. 114, 2331–2336. 10.1073/pnas.1621436114

Martinasso, G., Oraldi, M., Trombetta, A., Maggiora, M., Bertetto, O., Canuto, R.A., Muzio, G., 2007. Involvement of PPARs in Cell Proliferation and Apoptosis in Human Colon Cancer Specimens and in Normal and Cancer Cell Lines. PPAR Res. 2007, 93416. 10.1155/2007/93416

Massagué, J., 2012. TGFβ signalling in context. Nat. Rev. Mol. Cell Biol. 13, 616–630. 10.1038/nrm3434

McFadden, D.G., Barbosa, A.C., Richardson, J.A., Schneider, M.D., Srivastava, D., Olson, E.N., 2005. The Hand1 and Hand2 transcription factors regulate expansion of the embryonic cardiac ventricles in a gene dosage-dependent manner. Dev. Camb. Engl. 132, 189–201. 10.1242/dev.01562

McInnes, L., Healy, J., Melville, J., 2020. UMAP: Uniform Manifold Approximation and Projection for Dimension Reduction.

Melsted, P., Booeshaghi, A.S., Liu, L., Gao, F., Lu, L., Min, K.H.J., da Veiga Beltrame, E., Hjörleifsson, K.E., Gehring, J., Pachter, L., 2021. Modular, efficient and constant-memory single-cell RNA-seq preprocessing. Nat. Biotechnol. 39, 813–818. 10.1038/s41587-021-00870-2

Mommersteeg, M.T.M., Hoogaars, W.M.H., Prall, O.W.J., de Gier-de Vries, C., Wiese, C., Clout, D.E.W., Papaioannou, V.E., Brown, N.A., Harvey, R.P., Moorman, A.F.M., Christoffels, V.M., 2007. Molecular pathway for the localized formation of the sinoatrial node. Circ. Res. 100, 354–362. 10.1161/01.RES.0000258019.74591.b3

Moore, G.E., Ishida, M., Demetriou, C., Al-Olabi, L., Leon, L.J., Thomas, A.C., Abu-Amero, S., Frost, J.M., Stafford, J.L., Chaoqun, Y., Duncan, A.J., Baigel, R., Brimioulle, M., Iglesias-Platas, I., Apostolidou, S., Aggarwal, R., Whittaker, J.C., Syngelaki, A., Nicolaides, K.H., Regan, L., Monk, D., Stanier, P., 2015. The role and interaction of imprinted genes in human fetal growth. Philos. Trans. R. Soc. Lond. B. Biol. Sci. 370, 20140074. 10.1098/rstb.2014.0074

Moore, T., Haig, D., 1991. Genomic imprinting in mammalian development: a parental tug-of-war. Trends Genet. TIG 7, 45–49. 10.1016/0168-9525(91)90230-N

Moore-Morris, T., van Vliet, P.P., Andelfinger, G., Puceat, M., 2018. Role of Epigenetics in Cardiac Development and Congenital Diseases. Physiol. Rev. 98, 2453–2475. 10.1152/physrev.00048.2017

Morton, S.U., Brodsky, D., 2016. Fetal Physiology and the Transition to Extrauterine Life. Clin. Perinatol. 43, 395–407. 10.1016/j.clp.2016.04.001

Mullen, A.C., Orlando, D.A., Newman, J.J., Lovén, J., Kumar, R.M., Bilodeau, S., Reddy, J., Guenther, M.G., DeKoter, R.P., Young, R.A., 2011. Master transcription factors determine cell-type-specific responses to TGF-β signaling. Cell 147, 565–576. 10.1016/j.cell.2011.08.050

Murphy, S.A., Chen, E.Z., Tung, L., Boheler, K.R., Kwon, C., 2021a. Maturing heart muscle cells: Mechanisms and transcriptomic insights. Semin. Cell Dev. Biol. 119, 49–60. 10.1016/j.semcdb.2021.04.019

Murphy, S.A., Miyamoto, M., Kervadec, A., Kannan, S., Tampakakis, E., Kambhampati, S., Lin, B.L., Paek, S., Andersen, P., Lee, D.-I., Zhu, R., An, S.S., Kass, D.A., Uosaki, H., Colas, A.R., Kwon, C., 2021b. PGC1/PPAR drive cardiomyocyte maturation at single cell level via YAP1 and SF3B2. Nat. Commun. 12, 1648. 10.1038/s41467-021-21957-z

Naizhen, X., Kido, T., Yokoyama, S., Linnoila, R.I., Kimura, S., 2019. Spatiotemporal Expression of Three Secretoglobin Proteins, SCGB1A1, SCGB3A1, and SCGB3A2, in Mouse Airway Epithelia. J. Histochem. Cytochem. Off. J. Histochem. Soc. 67, 453–463. 10.1369/0022155419829050

Nestorowa, S., Hamey, F.K., Pijuan Sala, B., Diamanti, E., Shepherd, M., Laurenti, E., Wilson, N.K., Kent, D.G., Göttgens, B., 2016. A single-cell resolution map of mouse hematopoietic stem and progenitor cell differentiation. Blood 128, e20–31. 10.1182/blood-2016-05-716480

Olson, E.N., Schneider, M.D., 2003. Sizing up the heart: development redux in disease. Genes Dev. 17, 1937–1956. 10.1101/gad.1110103

Padula, S.L., Velayutham, N., Yutzey, K.E., 2021. Transcriptional Regulation of Postnatal Cardiomyocyte Maturation and Regeneration. Int. J. Mol. Sci. 22, 3288. 10.3390/ijms22063288

Paik, D.T., Cho, S., Tian, L., Chang, H.Y., Wu, J.C., 2020. Single-cell RNA sequencing in cardiovascular development, disease and medicine. Nat. Rev. Cardiol. 17, 457–473. 10.1038/s41569-020-0359-y

Park, K.-S., Rahat, B., Lee, H.C., Yu, Z.-X., Noeker, J., Mitra, A., Kean, C.M., Knutsen, R.H., Springer, D., Gebert, C.M., Kozel, B.A., Pfeifer, K., 2021. Cardiac pathologies in mouse loss of imprinting models are due to misexpression of H19 long noncoding RNA. eLife 10, e67250. 10.7554/eLife.67250

Pelengaris, S., Khan, M., Evan, G., 2002. c-MYC: more than just a matter of life and death. Nat. Rev. Cancer 2, 764–776. 10.1038/nrc904

Pervolaraki, E., Dachtler, J., Anderson, R.A., Holden, A.V., 2018. The developmental transcriptome of the human heart. Sci. Rep. 8, 15362. 10.1038/s41598-018-33837-6

Pinto, A.R., Ilinykh, A., Ivey, M.J., Kuwabara, J.T., D’Antoni, M.L., Debuque, R., Chandran, A., Wang, L., Arora, K., Rosenthal, N.A., Tallquist, M.D., 2016. Revisiting Cardiac Cellular Composition. Circ. Res. 118, 400–409. 10.1161/CIRCRESAHA.115.307778

Piquereau, J., Ventura-Clapier, R., 2018. Maturation of Cardiac Energy Metabolism During Perinatal Development. Front. Physiol. 9, 959. 10.3389/fphys.2018.00959

Purcell, N.H., Tang, G., Yu, C., Mercurio, F., DiDonato, J.A., Lin, A., 2001. Activation of NF-kappa B is required for hypertrophic growth of primary rat neonatal ventricular cardiomyocytes. Proc. Natl. Acad. Sci. U. S. A. 98, 6668–6673. 10.1073/pnas.111155798

Quaife-Ryan, G.A., Sim, C.B., Ziemann, M., Kaspi, A., Rafehi, H., Ramialison, M., El-Osta, A., Hudson, J.E., Porrello, E.R., 2017. Multicellular Transcriptional Analysis of Mammalian Heart Regeneration. Circulation 136, 1123–1139. 10.1161/CIRCULATIONAHA.117.028252

Rizzo, G., Gropper, J., Piollet, M., Vafadarnejad, E., Rizakou, A., Bandi, S.R., Arampatzi, P., Krammer, T., DiFabion, N., Dietrich, O., Arias-Loza, A.-P., Prinz, M., Mack, M., Schlepckow, K., Haass, C., Silvestre, J.-S., Zernecke, A., Saliba, A.-E., Cochain, C., 2023. Dynamics of monocyte-derived macrophage diversity in experimental myocardial infarction. Cardiovasc. Res. 119, 772–785. 10.1093/cvr/cvac113

Rudat, C., Kispert, A., 2012. Wt1 and epicardial fate mapping. Circ. Res. 111, 165–169. 10.1161/CIRCRESAHA.112.273946

Ruiz-Villalba, A., Romero, J.P., Hernández, S.C., Vilas-Zornoza, A., Fortelny, N., Castro-Labrador, L., San Martin-Uriz, P., Lorenzo-Vivas, E., García-Olloqui, P., Palacio, M., Gavira, J.J., Bastarrika, G., Janssens, S., Wu, M., Iglesias, E., Abizanda, G., de Morentin, X.M., Lasaga, M., Planell, N., Bock, C., Alignani, D., Medal, G., Prudovsky, I., Jin, Y.-R., Ryzhov, S., Yin, H., Pelacho, B., Gomez-Cabrero, D., Lindner, V., Lara-Astiaso, D., Prósper, F., 2020. Single-Cell RNA Sequencing Analysis Reveals a Crucial Role for CTHRC1 (Collagen Triple Helix Repeat Containing 1) Cardiac Fibroblasts After Myocardial Infarction. Circulation 142, 1831–1847. 10.1161/CIRCULATIONAHA.119.044557

Sakamoto, T., Matsuura, T.R., Wan, S., Ryba, D.M., Kim, J.U., Won, K.J., Lai, L., Petucci, C., Petrenko, N., Musunuru, K., Vega, R.B., Kelly, D.P., 2020. A Critical Role for Estrogen-Related Receptor Signaling in Cardiac Maturation. Circ. Res. 126, 1685–1702. 10.1161/CIRCRESAHA.119.316100

Schubert, M., Klinger, B., Klünemann, M., Sieber, A., Uhlitz, F., Sauer, S., Garnett, M.J., Blüthgen, N., Saez-Rodriguez, J., 2018. Perturbation-response genes reveal signaling footprints in cancer gene expression. Nat. Commun. 9, 20. 10.1038/s41467-017-02391-6

Seidman, C.E., Schmidt, E.V., Seidman, J.G., 1991. cis-dominance of rat atrial natriuretic factor gene regulatory sequences in transgenic mice. Can. J. Physiol. Pharmacol. 69, 1486–1492. 10.1139/y91-223

Siedner, S., Krüger, M., Schroeter, M., Metzler, D., Roell, W., Fleischmann, B.K., Hescheler, J., Pfitzer, G., Stehle, R., 2003. Developmental changes in contractility and sarcomeric proteins from the early embryonic to the adult stage in the mouse heart. J. Physiol. 548, 493–505. 10.1113/jphysiol.2002.036509

Sim, C.B., Ziemann, M., Kaspi, A., Harikrishnan, K.N., Ooi, J., Khurana, I., Chang, L., Hudson, J.E., El-Osta, A., Porrello, E.R., 2015. Dynamic changes in the cardiac methylome during postnatal development. FASEB J. Off. Publ. Fed. Am. Soc. Exp. Biol. 29, 1329–1343. 10.1096/fj.14-264093

Skelly, D.A., Squiers, G.T., McLellan, M.A., Bolisetty, M.T., Robson, P., Rosenthal, N.A., Pinto, A.R., 2018. Single-Cell Transcriptional Profiling Reveals Cellular Diversity and Intercommunication in the Mouse Heart. Cell Rep. 22, 600–610. 10.1016/j.celrep.2017.12.072

Sparago, A., Cerrato, F., Vernucci, M., Ferrero, G.B., Silengo, M.C., Riccio, A., 2004. Microdeletions in the human H19 DMR result in loss of IGF2 imprinting and Beckwith-Wiedemann syndrome. Nat. Genet. 36, 958–960. 10.1038/ng1410

Subramanian, A., Tamayo, P., Mootha, V.K., Mukherjee, S., Ebert, B.L., Gillette, M.A., Paulovich, A., Pomeroy, S.L., Golub, T.R., Lander, E.S., Mesirov, J.P., 2005. Gene set enrichment analysis: a knowledge-based approach for interpreting genome-wide expression profiles. Proc. Natl. Acad. Sci. U. S. A. 102, 15545–15550. 10.1073/pnas.0506580102

Sun, Y., Liang, X., Najafi, N., Cass, M., Lin, L., Cai, C.-L., Chen, J., Evans, S.M., 2007. Islet 1 is expressed in distinct cardiovascular lineages, including pacemaker and coronary vascular cells. Dev. Biol. 304, 286–296. 10.1016/j.ydbio.2006.12.048

Suo, S., Zhu, Q., Saadatpour, A., Fei, L., Guo, G., Yuan, G.-C., 2018. Revealing the Critical Regulators of Cell Identity in the Mouse Cell Atlas. Cell Rep. 25, 1436–1445.e3. 10.1016/j.celrep.2018.10.045

Szabó, P.E., Tang, S.-H.E., Silva, F.J., Tsark, W.M.K., Mann, J.R., 2004. Role of CTCF binding sites in the Igf2/H19 imprinting control region. Mol. Cell. Biol. 24, 4791–4800. 10.1128/MCB.24.11.4791-4800.2004

Talman, V., Teppo, J., Pöhö, P., Movahedi, P., Vaikkinen, A., Karhu, S.T., Trošt, K., Suvitaival, T., Heikkonen, J., Pahikkala, T., Kotiaho, T., Kostiainen, R., Varjosalo, M., Ruskoaho, H., 2018. Molecular Atlas of Postnatal Mouse Heart Development. J. Am. Heart Assoc. 7, e010378. 10.1161/JAHA.118.010378

Tan, C.M.J., Lewandowski, A.J., 2020. The Transitional Heart: From Early Embryonic and Fetal Development to Neonatal Life. Fetal Diagn. Ther. 47, 373–386. 10.1159/000501906

Thomas, A.A., Biswas, S., Feng, B., Chen, S., Gonder, J., Chakrabarti, S., 2019. lncRNA H19 prevents endothelial-mesenchymal transition in diabetic retinopathy. Diabetologia 62, 517–530. 10.1007/s00125-018-4797-6

Tombor, L.S., John, D., Glaser, S.F., Luxán, G., Forte, E., Furtado, M., Rosenthal, N., Baumgarten, N., Schulz, M.H., Wittig, J., Rogg, E.-M., Manavski, Y., Fischer, A., Muhly-Reinholz, M., Klee, K., Looso, M., Selignow, C., Acker, T., Bibli, S.-I., Fleming, I., Patrick, R., Harvey, R.P., Abplanalp, W.T., Dimmeler, S., 2021. Single cell sequencing reveals endothelial plasticity with transient mesenchymal activation after myocardial infarction. Nat. Commun. 12, 681. 10.1038/s41467-021-20905-1

Tucker, N.R., Chaffin, M., Fleming, S.J., Hall, A.W., Parsons, V.A., Bedi, K.C., Akkad, A.-D., Herndon, C.N., Arduini, A., Papangeli, I., Roselli, C., Aguet, F., Choi, S.H., Ardlie, K.G., Babadi, M., Margulies, K.B., Stegmann, C.M., Ellinor, P.T., 2020. Transcriptional and Cellular Diversity of the Human Heart. Circulation 142, 466–482. 10.1161/CIRCULATIONAHA.119.045401

Uosaki, H., Cahan, P., Lee, D.I., Wang, S., Miyamoto, M., Fernandez, L., Kass, D.A., Kwon, C., 2015. Transcriptional Landscape of Cardiomyocyte Maturation. Cell Rep. 13, 1705–1716. 10.1016/j.celrep.2015.10.032

Uscategui Calderon, M., Gonzalez, B.A., Yutzey, K.E., 2023. Cardiomyocyte-fibroblast crosstalk in the postnatal heart. Front. Cell Dev. Biol. 11, 1163331. 10.3389/fcell.2023.1163331

Vafadarnejad, E., Rizzo, G., Krampert, L., Arampatzi, P., Arias-Loza, A.-P., Nazzal, Y., Rizakou, A., Knochenhauer, T., Bandi, S.R., Nugroho, V.A., Schulz, D.J.J., Roesch, M., Alayrac, P., Vilar, J., Silvestre, J.-S., Zernecke, A., Saliba, A.-E., Cochain, C., 2020. Dynamics of Cardiac Neutrophil Diversity in Murine Myocardial Infarction. Circ. Res. 127, e232–e249. 10.1161/CIRCRESAHA.120.317200

Van de Sande, B., Flerin, C., Davie, K., De Waegeneer, M., Hulselmans, G., Aibar, S., Seurinck, R., Saelens, W., Cannoodt, R., Rouchon, Q., Verbeiren, T., De Maeyer, D., Reumers, J., Saeys, Y., Aerts, S., 2020. A scalable SCENIC workflow for single-cell gene regulatory network analysis. Nat. Protoc. 15, 2247–2276. 10.1038/s41596-020-0336-2

Vanlandewijck, M., He, L., Mäe, M.A., Andrae, J., Ando, K., Del Gaudio, F., Nahar, K., Lebouvier, T., Laviña, B., Gouveia, L., Sun, Y., Raschperger, E., Räsänen, M., Zarb, Y., Mochizuki, N., Keller, A., Lendahl, U., Betsholtz, C., 2018. A molecular atlas of cell types and zonation in the brain vasculature. Nature 554, 475–480. 10.1038/nature25739

Viereck, J., Bührke, A., Foinquinos, A., Chatterjee, S., Kleeberger, J.A., Xiao, K., Janssen-Peters, H., Batkai, S., Ramanujam, D., Kraft, T., Cebotari, S., Gueler, F., Beyer, A.M., Schmitz, J., Bräsen, J.H., Schmitto, J.D., Gyöngyösi, M., Löser, A., Hirt, M.N., Eschenhagen, T., Engelhardt, S., Bär, C., Thum, T., 2020. Targeting muscle-enriched long non-coding RNA H19 reverses pathological cardiac hypertrophy. Eur. Heart J. 41, 3462–3474. 10.1093/eurheartj/ehaa519

Wang, Y., Yao, F., Wang, Lipeng, Li, Z., Ren, Z., Li, D., Zhang, M., Han, L., Wang, S.-Q., Zhou, B., Wang, Li, 2020. Single-cell analysis of murine fibroblasts identifies neonatal to adult switching that regulates cardiomyocyte maturation. Nat. Commun. 11, 2585. 10.1038/s41467-020-16204-w

Wu, B., Wang, Y., Lui, W., Langworthy, M., Tompkins, K.L., Hatzopoulos, A.K., Baldwin, H.S., Zhou, B., 2011. Nfatc1 coordinates valve endocardial cell lineage development required for heart valve formation. Circ. Res. 109, 183–192. 10.1161/CIRCRESAHA.111.245035

Wünnemann, F., Ta-Shma, A., Preuss, C., Leclerc, S., van Vliet, P.P., Oneglia, A., Thibeault, M., Nordquist, E., Lincoln, J., Scharfenberg, F., Becker-Pauly, C., Hofmann, P., Hoff, K., Audain, E., Kramer, H.-H., Makalowski, W., Nir, A., Gerety, S.S., Hurles, M., Comes, J., Fournier, A., Osinska, H., Robins, J., Pucéat, M., MIBAVA Leducq Consortium principal investigators, Elpeleg, O., Hitz, M.-P., Andelfinger, G., 2020. Loss of ADAMTS19 causes progressive non-syndromic heart valve disease. Nat. Genet. 52, 40–47. 10.1038/s41588-019-0536-2

Xie, Z., Bailey, A., Kuleshov, M.V., Clarke, D.J.B., Evangelista, J.E., Jenkins, S.L., Lachmann, A., Wojciechowicz, M.L., Kropiwnicki, E., Jagodnik, K.M., Jeon, M., Ma’ayan, A., 2021. Gene Set Knowledge Discovery with Enrichr. Curr. Protoc. 1, e90. 10.1002/cpz1.90

Xiong, L., Sun, Y., Huang, J., Ma, P., Wang, X., Wang, J., Chen, B., Chen, J., Huang, M., Huang, S., Liu, Y., 2022. Long Non-Coding RNA H19 Prevents Lens Fibrosis through Maintaining Lens Epithelial Cell Phenotypes. Cells 11, 2559. 10.3390/cells11162559

Yamada, S., Nomura, S., 2020. Review of Single-Cell RNA Sequencing in the Heart. Int. J. Mol. Sci. 21, 8345. 10.3390/ijms21218345

Zhang, H., Pu, W., Li, G., Huang, X., He, L., Tian, X., Liu, Q., Zhang, L., Wu, S.M., Sucov, H.M., Zhou, B., 2016. Endocardium Minimally Contributes to Coronary Endothelium in the Embryonic Ventricular Free Walls. Circ. Res. 118, 1880–1893. 10.1161/CIRCRESAHA.116.308749

Zhang, T., Liu, J., Zhang, J., Thekkethottiyil, E.B., Macatee, T.L., Ismat, F.A., Wang, F., Stoller, J.Z., 2013. Jun is required in Isl1-expressing progenitor cells for cardiovascular development. PloS One 8, e57032. 10.1371/journal.pone.0057032

Zhao, M.-T., Ye, S., Su, J., Garg, V., 2020. Cardiomyocyte Proliferation and Maturation: Two Sides of the Same Coin for Heart Regeneration. Front. Cell Dev. Biol. 8, 594226. 10.3389/fcell.2020.594226

Zhou, B., Wu, B., Tompkins, K.L., Boyer, K.L., Grindley, J.C., Baldwin, H.S., 2005. Characterization of Nfatc1 regulation identifies an enhancer required for gene expression that is specific to pro-valve endocardial cells in the developing heart. Dev. Camb. Engl. 132, 1137–1146. 10.1242/dev.01640

Zhou, P., Pu, W.T., 2016. Recounting Cardiac Cellular Composition. Circ. Res. 118, 368–370. 10.1161/CIRCRESAHA.116.308139

